# 5-Iodotubercidin inhibits Epithelial to Mesenchymal Transition by inhibiting IKK/NFκB-dependent gene expression

**DOI:** 10.64898/2026.02.13.705677

**Authors:** Sayari Bhattacharya, Muskan Manjari, Vaishnavi Ganesh, Manoj B. Menon, Sonam Dhamija

## Abstract

Epithelial to mesenchymal transition (EMT) is a process of trans-differentiation important for development, inflammation and cancer. Transforming Growth Factor-β (TGFβ) is a physiologically relevant inducer of EMT. We had recently characterized the adenosine analogue, adenosine kinase inhibitor 5-Iodotubercidin (5-ITu), as a compound which preferentially sensitizes MK2-deficient cells to TNF-induced, RIPK1-dependent cell death. Here we investigated the effect of 5-ITu on TGFβ-induced EMT. 5-ITu suppressed TGFβ-induced morphological changes and migration in A549 (lung cancer) and PANC1 (pancreatic cancer) cell lines. Consistent with these effects, there was significant suppression of EMT markers as indicated by qPCR, immunoblotting and immunofluorescence and confocal microscopy. Mechanistic investigations revealed that 5-ITu-mediated EMT suppression was independent of adenosine kinase inhibition and RIPK1 activation. 5-ITu suppressed NFκB activity in cells undergoing EMT and IKK inhibition phenocopied the effect of 5-ITu on EMT. Kinase assays revealed IKKβ as a potential direct target of 5-ITu. We identified a TGFβ-associated, NFκB-dependent gene signature consisting of 4 genes, which are differentially regulated upon 5-ITu treatment. Interestingly, this 4 gene signature could predict survival in lung and pancreatic cancer. The identification of this role for the multitarget kinase inhibitor 5-ITu in NFκB activity-dependent EMT, in addition to RIPK1-dependent necroptosis has potential implications in anticancer strategies.

## Introduction

Epithelial to Mesenchymal Transition (EMT) is a trans-differentiation process where epithelial cells lose their inherent features like cell-cell contacts, cell polarity and gain mesenchymal phenotypes (1). EMT is a critical player in development and in the process of cancer progression and metastasis. TGFβ is the best characterized cytokine known to induce EMT in a wide variety of mammalian epithelial cells and cell line models (2, 3). Downstream to TGFβ-receptors, SMAD2/3/4 transcription factors regulate key players that orchestrate the EMT program. The canonical EMT signaling includes the repression of epithelial genes like E-cadherin (*CDH1*) and the upregulation of mesenchymal genes like Vimentin (*VIM*), N-cadherin (*CDH2*), and fibronectin (*FN1*). SMAD signaling mediates the transcriptional reprogramming by inducing the expression of key transcriptional factors such as Snail (SNAI1), Slug (SNAI2), ZEB1, and ZEB2, which in turn suppress CDH1 expression (4). Concurrently, they promote the transcription of mesenchymal genes conferring migratory and invasive properties. In addition, other non-canonical signaling pathways activated by TGFβ, and involved in EMT include MAPK pathways, PI3K/Akt pathway, NFκB and Rho-like GTPase signaling (3, 5).

5-Iodotubercidin (5-ITu) is a nucleoside analogue and has been reported as a potent adenosine kinase inhibitor (6). *In vitro* and *in vivo* studies have established strong anti-cancer potential of 5-ITu (7). Adenosine kinase inhibition by 5-ITu was shown to suppress SARS-CoV2 replication (8). Despite the discovery as an adenosine kinase inhibitor, 5-ITu was later characterized as a protein kinase inhibitor (9). The earliest characterised protein kinase targets of 5-ITu include CK1, CK2 and ERK2 (10, 11). In addition, DYRK1A inhibition by 5-ITu was shown to modulate β-cell proliferation and function (12). Another well-established target of 5-ITu is the histone-H3 kinase Haspin/GSG2(13-15).

In our recent study, we identified 5-ITu as a small molecule modulator of RIPK1-dependent death. In a small molecule inhibitor screen, 5-ITu was shown to specifically sensitize MK2-deficient cells to TNF-induced necroptosis (16). TNF-induced cell death is modulated mainly by two checkpoints involving IKK and MK2-mediated RIPK1 phosphorylation, which prevents RIPK1 autophosphorylation and subsequent cytotoxic response (17-21). Mechanistically, 5-ITu mediated inhibition of IKK-dependent RIPK1 phosphorylation was found to facilitate RIPK1 autophosphorylation and predominant necroptotic response in the cells lacking MK2 activity (16).

In the present work, we investigated the effect of 5-ITu on EMT. 5-ITu strongly suppressed TGFβ-induced EMT as evident from EMT marker expression, morphological analyses and cell migration. We present evidence that 5-ITu inhibits NFκB signaling, by directly targeting the IKKs, independent of adenosine kinase inhibition. In contrast to the effect of 5-ITu on TNF-induced cytotoxicity, the suppression of TGFβ-induced EMT was independent of RIPK1 activity. Moreover, we identified an NFκB-dependent gene signature, sensitive to 5-ITu, and modulated by TGFβ. Interestingly, TCGA (The Cancer Genome Atlas) data analyses revealed that this 4-gene signature is associated with poor survival in lung and pancreatic cancer patients.

## Materials and Methods

Cell culture and treatments: PANC1 and HEK293T cells were cultured in DMEM High glucose media (#AL007A, HiMedia) and A549 cells were maintained in DMEM/F12 (#AL215A, HiMedia,) supplemented with 10% fetal calf serum (FCS) and 1x antibiotic-antimycotic solution (#15240062, Gibco) at 37⍰°C, in a humidified atmosphere supplemented with 5% CO_2_. ⍰ Cells were treated as indicated in 10 cm dishes (cell-cycle analyses), 12 well plates (for immunoblot analyses), 6 well plates (RNA isolation), 24 well plates (reporter assays) or 96 well plates (viability, cytotoxicity and wound-healing assays). Cells were pretreated with inhibitors or DMSO as solvent control before TNFα and/or TGFβ stimulation wherever indicated. All treatments were performed in normal growth medium for the respective cell lines.

### Antibodies and reagents

N-Cadherin/CDH2 (#13116), E-Cadherin/CDH1 (#3195), SNAIL/SNAI1 (#3879), ZEB1 (#3396), Vimentin (#5741), pS^536^-p65 (#3033), NF-κB p65 (#8242), IKKβ (#8943) and pS^32/36^-IkBα (#9246) antibodies were from Cell Signaling Technology (CST). Further antibodies used were against IkBα (#AB0040, Bio Bharati LifeScience), GAPDH (#MAB932Hu23, Cloud-Clone Corp.), α Tubulin (#12G10, DSHB), GST (#sc-138, Santa Cruz Biotechnology). Goat anti-mouse IgG (H+L)-HRP (#115-035-003), and goat anti-rabbit IgG (H+L)-HRP (111-035-045) secondary antibodies were from Jackson Immuno Research Laboratories. Alexa-488 Goat anti-Rabbit IgG (H+L) antibody was from Elabscience (#E-AB-1055) and Phalloidin iFluor™ 647 Conjugate (#20555, Cayman chemicals). Other chemicals and reagents used were: DAPI (#80145, Sisco Research Laboratories), ATP (#RM439, HiMedia), DiR (DiIC18) (#60017, Biotium Inc.).

The source and working concentrations used for the stimulants and inhibitors were: recombinant human TGFβ (#100-21-10, Peprotech, 5 ng/mL) and recombinant human TNFα (#PHC3016, Gibco, 101ng/ml), 5-Iodotubercidin (#HY-15424, 0.1-10 µM), IKKα/β inhibitor BMS345541 Hydrochloride (#HY-10518, 5 µM) (IKKβ IC_50_= 0.3 μM, IKKα IC_50_= 4 μM)(22), RIPK1 inhibitor Nec-1 (#HY-15760, 50 µM), GSK481 (HY-100131, 5 µM), ABT-702 dihydrochloride (#HY-103161, 5-10 µM) were obtained from MedChemExpress.

### Analysis of cell viability and cell death

A549 and PANC1 cells were seeded in triplicates in 96 well plates in 100 µL media per well. The cell numbers were optimized as 6000 cells/well for both the cell lines for 48h treatment. Next day, cells were treated with various indicated combinations of reagents. For measuring cell death by Sytox Green assay (#S7020, Invitrogen, 5 mM stock in DMSO), Sytox green solution was also added during the time of treatments at a final concentration of 0.25 µM and fluorescence measurements were made using a multimode reader (Cytation 5, BioTek). Cell viability was evaluated using WST-8 (#72465, SRL) reagent (23). A549 and PANC1 cells were seeded in triplicates in 96 well plates in 100 µL media per well. Next day, cells were treated with various combinations of reagents as indicated for 48 h. WST-8 (10 µL/well) was added directly to the wells in complete medium and plates were then incubated in a cell culture incubator for 45 – 60 min at 37°C and the absorbance was measured at 450 nm using a microplate reader (Cytation 5, BioTek).

### Quantitative scratch-would healing assays

Fluorescence-based scratch wound healing assay were performed as described previously (24). In brief, DiR-labeled A549 or PANC1 cells were seeded in 96 well plates at a density of 40000 cells per well. After 18h of seeding, scratches were made using 10 µl tips. At the beginning of experiment (t=0) and after allowing migration for 18, or 24 h, images were acquired using the 800 nm channel (42 µm resolution) using a near-infrared fluorescence scanner (Odyssey DLx, LICORbio). Scans were converted to 8-bit images and analyzed using ImageJ and data is represented as migration indices.

### Cell cycle analyses

A549 cells (9 x 10^5^ cells) were seeded in 10 cm dishes and were treated with TGF-β and/or 5-ITu (5 µM) as indicated for 24 h. The cells were harvested, washed with 1x phosphate-buffered saline (PBS), and fixed in ice-cold 90 % ethanol. After fixation, cells were washed with 1x PBS, resuspended in the staining solution (100 µg/mL RNase A and 100 µg/mL Propidium Iodide in 1x PBS), and incubated in the dark for 30 minutes. DNA content was analyzed using a BD Accuri C6 Plus flow cytometer at low flow rate, and cell cycle distribution plotted and quantified using the Floreada.io software (https://floreada.io, accessed in August 2025).

### NFκB-luciferase reporter assays

HEK293T cells were transfected using polyethylenimine reagent (#408727, Sigma Aldrich) with reporter plasmids followed by media change after 6 h. A 3× κB luciferase reporter construct (encoding firefly luciferase, a gift from Dr. Mark Windheim, Hannover Medical School) (25) and pGL4.70-Renilla luciferase (normalisation control, Promega) were used. Eighteen hours post transfection cells were treated with TNFα or TGFβ along with 5-ITu (5 µM) and BMS345541 (5 µM) for additional six hours. The plates were lysed twenty four hours post transfection in 1x Passive Lysis Buffer (#E1941, Promega). Reporter assays were performed using in house buffers as reported previously (26) and luminescence measured using a multi-mode reader (Cytation 5, BioTek).

### *In vitro* kinase assays

For *In vitro* kinase assays, 20 ng recombinant GST IKKβ (#PV3836, Invitrogen) and 1 µg recombinant IκBα (#BB-SPP40, Bio Bharati LifeScience) were used in a reaction volume of 25 µL. The kinase was incubated with inhibitors at indicated concentrations at 30°C for 5 mins followed by addition of substrate, and ATP mix, and were incubated for 30 min at 30 °C. The final reaction buffer consisted of 20 mM HEPES, 20 mM β-glycerophosphate, 10 mM NaF, 10 mM MgCl2 and 25 µM ATP. Reaction was stopped by adding 5X SDS loading dye and monitored by immunoblotting.

### RNA isolation, cDNA synthesis and qPCR analyses

A549 and PANC1 cells were treated for 24 h in 6 well plates, total RNA was isolated using the NucleoSpin RNA extraction kit (Macherey & Nagel) and cDNA was synthesized using a cDNA synthesis kit (BB-E0043, Bio Bharati LifeScience,) using random hexamer primers according to the manufacturer’s instructions. Real time PCR quantitation was performed using TB Green Premix Ex Taq II (Tli RNase H Plus, Takara) on a CFX96 system (Bio-Rad). Gene expression was normalized to *GAPDH* or Cyclophilin/*PPIA*. The primer sequences used are listed in the Supplementary table S1.

### Western blotting

After 24h treatments with TGFβ and/or inhibitors, cells were lysed in kinase-assay lysis buffer (20 mM Tris, 1 mM Na_3_VO_4_, 1 mM EDTA, 1 mM EGTA, 10 mM β-glycerophosphate, 50 mM NaF, 1% Triton X-100) supplemented with protease inhibitor cocktail (#11873580001,Sigma Aldrich, and phosphatase inhibitor cocktails (#04906845001,Phosstop, Roche), and total protein quantified by Bradford assay (#5000205, Bio-Rad) before processing for immunoblotting. SDS-PAGE (10 %) gels were used for protein separation followed by western blotting using nitrocellulose membrane (#10600001, Bio-Rad). Membranes were blocked with 5 % powdered skim milk in PBS with 0.1 % Tween 20 for 1h at room temperature (RT), followed by incubation with the primary antibodies at 4 °C overnight. Blots were washed, incubated with secondary antibodies for one hour at RT and were developed using ECL reagents (Westarnova 2.0, Cyanagen; Clarity Max, Bio-Rad) and bands were visualized using Fusion Solo 6S chemiluminescence imager (Vilber).

### Immunostaining and Microscopy

For obtaining phase contrast images, A549 and PANC1 cells were seeded in 12 well plates and treated with TGFβ only and co-treated with 5-ITu and SB42 or with 5-ITu only for 24h. Live phase-contrast images were captured using camera attached to the microscope (#PZQ-106S, Quasmo, India). For Immuno-fluorescence staining, cells were seeded on poly-L-lysine precoated glass cover slips and processed as reported previously (27). Primary antibodies against CDH1 were used at a 1:200 dilution in 1% BSA–PBS for 2 h at RT. Alexa-488 labelled secondary antibodies and fluorescent labeled Phalloidin were used at 1:250 dilution in 1% BSA-PBS at RT for 45 min, followed by DAPI (500 ng/mL in PBS) staining for 10 min at RT. Imaging was performed using a Leica TCS SP8 confocal microscope with standard settings.

### *In silico* data analyses

HALLMARK_TGF_BETA_SIGNALING and HALLMARK_TNFA_SIGNALING_VIA_NFKB genesets were obtained from the Molecular Signature Database (www.gsea-msigdb.org)(28, 29). Venn diagrams were created using Venny 2.1 (https://bioinfogp.cnb.csic.es/tools/venny/)(30). We used the GEPIA2 server (http://gepia2.cancer-pku.cn/) (31) to access TCGA (https://www.cancer.gov/tcga) data and perform survival analyses (overall survival and disease-free survival) of TCGA lung cancer (lung adenocarcinoma and squamous cell carcinoma) and pancreatic cancer data.

### Statistics and Reproducibility

All cell death and viability assays were performed in biological triplicate samples and the data presented are representative of three independent experiments. All Immunoblot results presented in figures are representative results from at least three independent experiments, unless indicated otherwise in the figure legends. Migration assays were performed with a minimum of four biological replicates and the data presented are representative of at least two independent experiments. The calculations, statistical analyses and graphs were performed using Microsoft Excel. Two-tailed unpaired *t*-test was used to calculate statistical significance for quantitative assays. The statistics and source data are presented in Supplementary Table S2.

## Results

### 5-ITu inhibits TGFβ-induced epithelial to mesenchymal transition

Our previous studies have identified 5-ITu as an inhibitor of TNF-induced survival signaling and 5-ITu was shown to facilitate TNF-induced cell death synergistically with p38/MK2 inactivation (16). To understand the effect of 5-ITu on other physiologically and pathologically relevant events, we focused on TGFβ-induced epithelial to mesenchymal transition (EMT). Co-treatment with 5-ITu suppressed the TGFβ-induced morphological changes in A549 (lung adenocarcinoma) and PANC1 (pancreatic ductal carcinoma) cell lines (Figure 1A & Supplementary Figure S1A). Consistent with the EMT-like morphological changes, immunofluorescence analyses revealed upregulation of filamentous actin (phalloidin staining) and re-distribution of epithelial marker E-cadherin (CDH1) in TGFβ-treated A549 cells. These effects were also suppressed by treatment with 5-ITu (Figure 1B). To quantitatively assess the effect of 5-ITu on TGFβ-induced EMT in A549 cells, we monitored the expression of epithelial (E-cadherin/*CDH1*) and mesenchymal (N-cadherin/*CDH2, ZEB1*, Snail 1/*SNAI1*) marker genes by realtime qPCR. 5-ITu co-treatment significantly suppressed the TGFβ-induced upregulation of mesenchymal transcripts and down regulation of epithelial marker *CDH1* (Figure 1C). While the effects of 5-ITu on TGFβ-regulated marker gene expression were comparable to that of the TGF-receptor kinase inhibitor SB431542, 5-ITu additionally suppressed the basal expression levels of EMT transcription factors *SNAI1* and *ZEB1*. Similar results were also observed in TGFβ-treated PANC1 cells (Supplementary Figure S1B). Consistent with the effect at RNA level, TGFβ-induced changes in CDH1, CDH2and SNAI1 protein levels were also reversed by 5-ITu treatment (Figure 1D). EMT is associated with enhanced capacity for cell migration often evaluated by scratch wound-healing assays (24). TGFβ-treatment increased the wound closure capacity of both A549 and PANC1 cells, when assayed by a quantitative scratch wound healing assay. 5-ITu dose-dependently inhibited the wound closure in both cell lines (Figure 1E). Consistent with the effects on mesenchymal marker expression, 5-ITu mediated suppression of migration was stronger than that of SB431542. These findings clearly establish 5-ITu as an inhibitor of EMT.

**Figure 1.**
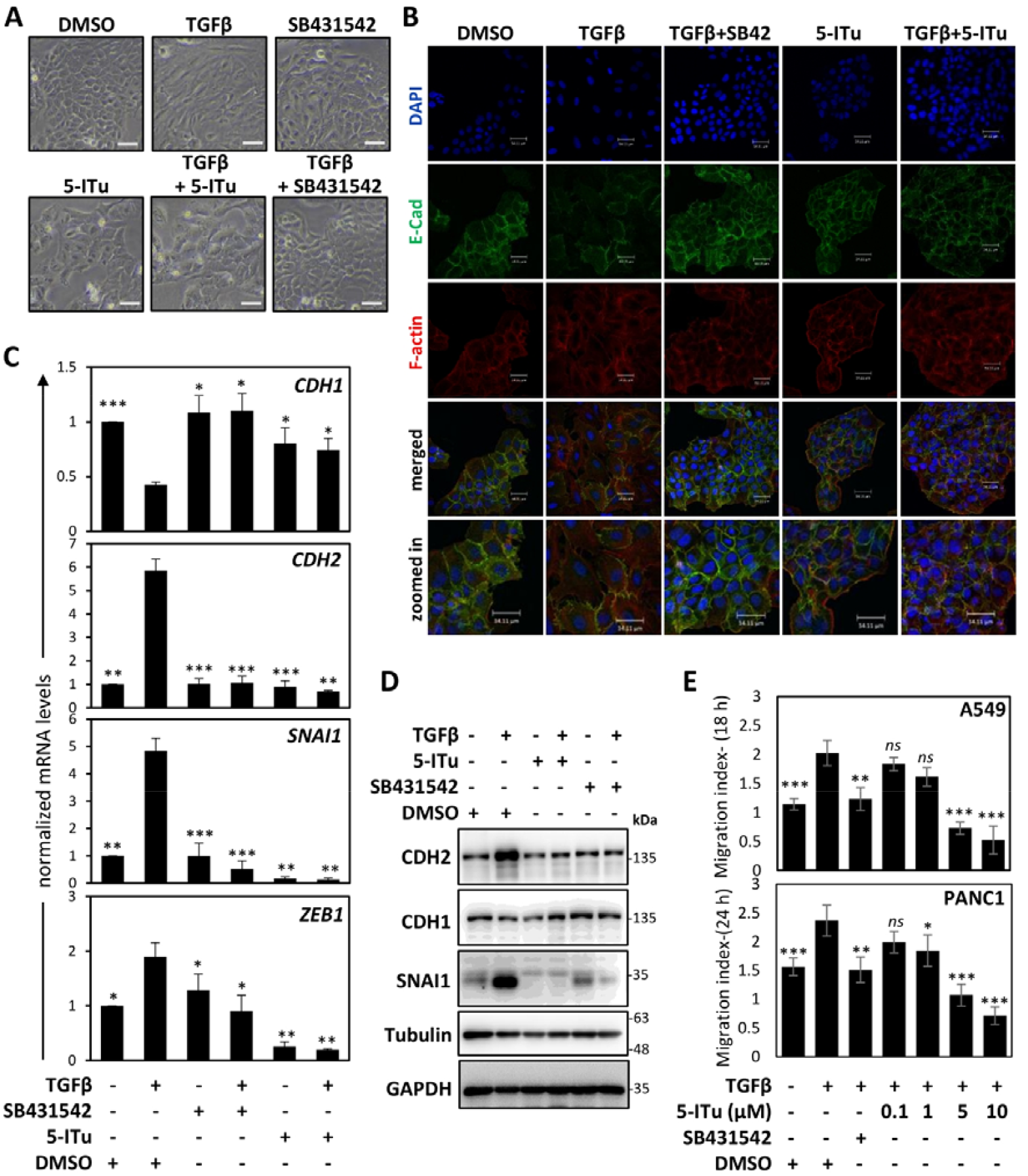
5-Iodotubercidin (5-ITu) inhibit TGFβ-induced EMT. **A**. A549 cells were treated as indicated for 24 h and images were acquired using a phase-contrast 10x objective. Scale bars indicate 50 µm. **B**. Cells seeded on cover slips were treated for 24 h as indicated. After fixation, cells were stained for F-actin (phalloidin), DNA (DAPI) and E-cadherin/CDH1 (E-cad) and were imaged by confocal microscopy. **C & D**. A549 Cells were stimulated with TGFβ in combination with indicated inhibitors or solvent control (DMSO) and EMT marker expression was monitored by qPCR (n=3, mean±SD for 3 independent experiments) (C) and immunoblotting (**D**). **E**. A549 (upper panel) and (PANC1) cells were subjected to a quantitative scratch wound healing assay for 18 h and 24 h respectively and migration indices were plotted (n≥4, mean±SD). (* denotes p-value ≤ 0.05, ** denotes p-value ≤ 0.01, *** denotes p-value ≤ 0.001).

### Adenosine kinase inhibition does not phenocopy the effect of 5-ITu on EMT

5-ITu is a nucleoside analogue and is best characterised as an adenosine kinase inhibitor (32). Adenosine kinase inhibition can lead to the accumulation of extracellular adenosine driving purinergic signaling, with potential effect on EMT. To understand whether 5-ITu-mediated suppression is an after effect of adenosine kinase inhibition, we monitored the effect of a structurally unrelated adenosine kinase inhibitor ABT-702 on TGFβ-induced EMT (33). Unlike 5-ITu, ABT-702-treatment did not significantly alter the TGFβ-induced morphological changes in A549 and PANC1 cells (Figure 2A). There was a minor, but significant effect of ABT-702 on TGFβ-induced expression of SNAI1 in A549 cells, but similar effects were not observed on the expression of other marker genes tested (Figure 2B & 2D). Comparable effects were observed in PANC1 cells (Figure 2C & 2E) with no inhibition of TGFβ-induced EMT markers seen with ABT-702. While there was significant inhibitory effect of ABT-702 on TGFβ-induced migration in A549 cells, the effect was not observed in PANC1 cells (Figure 2F & 2G). Moreover, the effect of 5 µM 5-ITu was significantly stronger than the effect shown by ABT-702 in either cell lines. These results indicate that the EMT-inhibitory effect of 5-ITu is probably independent of its effect on adenosine kinase activity.

**Figure 2.**
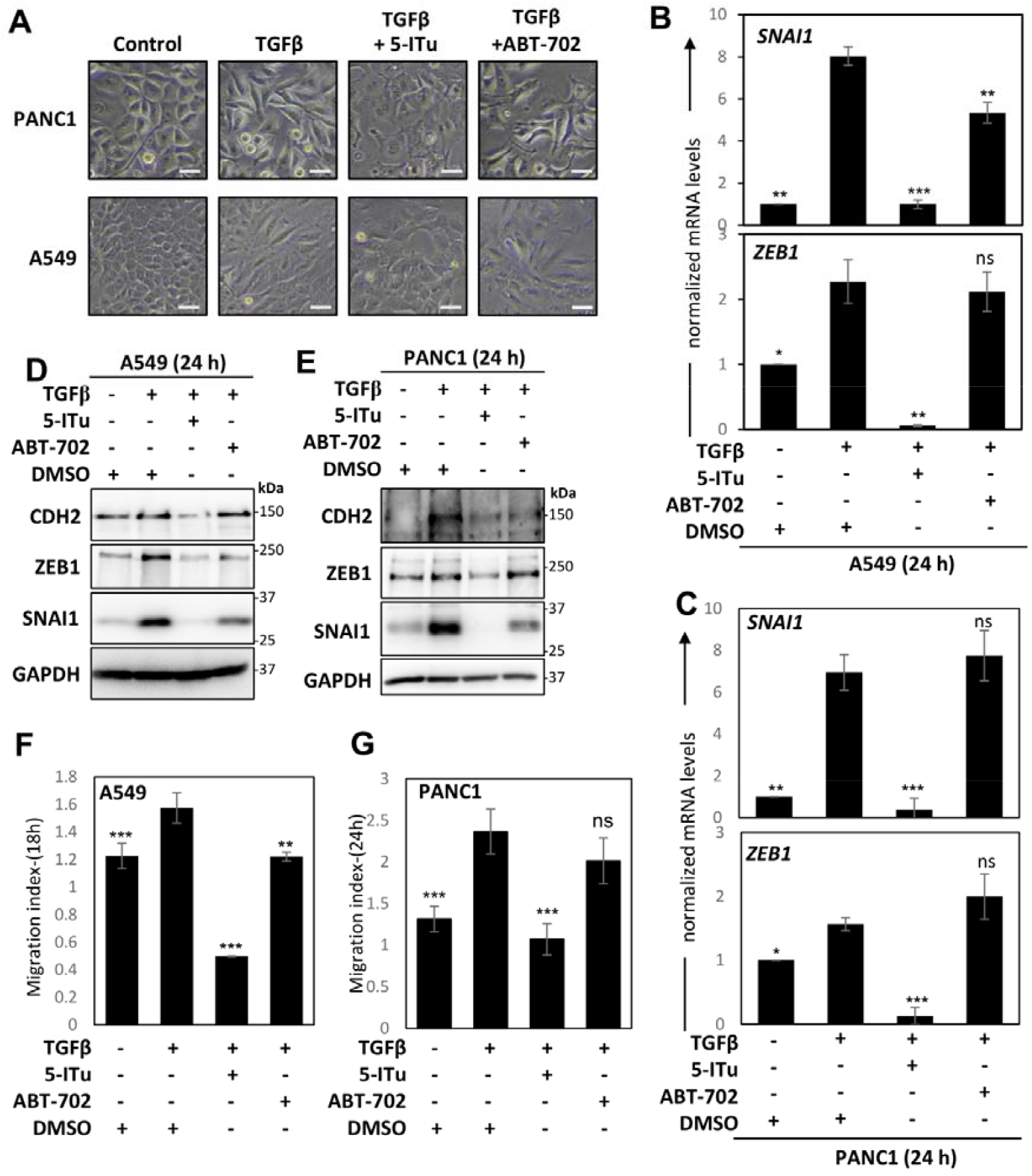
Adenosine kinase inhibition does not phenocopy the effect of 5-ITu on EMT. **A** A549 cells were treated as indicated for 24 h and images were acquired using a phase-contrast 10x objective. Scale bars indicate 50µm. **B - E**. A549 (B & D) or PANC1 (C & E) cells were stimulated with TGFβ in combination with indicated inhibitors or solvent control (DMSO) and EMT marker expression was monitored by qPCR (n=3, mean±SD plotted) (B & C) and immunoblotting (D & E). F & G. A549 (F) or PANC1 (G) cells were subjected to a quantitative scratch wound healing assay in the presence of stimulants/inhibitors as indicated and migration indices were plotted (n≥5, mean±SD). (* denotes p-value ≤ 0.05, ** denotes p-value ≤ 0.01, *** denotes p-value ≤ 0.001).

### The effect of 5-ITu on EMT is independent of RIPK1-mediated cytotoxicity

Our previous work had established an adenosine-kinase independent role for 5-ITu in facilitating TNF and LPS-induced apoptosis/necroptosis through the hyperactivation of RIPK1. To monitor whether a similar cytotoxic response is responsible for the effect of 5-ITu on TGFβ-induced EMT, we performed cell viability assays comparing the effect of 5-ITu on TGFβ and TNFα treatment. When cell viability was assayed using WST-8 assays, 5-ITu displayed a dose-dependent decrease in cell viability at 48 h. However, there was no effect of TGFβ co-treatment on viability. On the other hand, co-treatment with TNFα induced further loss of A549 cell viability (Figure 3A). We also monitored the presence of cell death in TGFβ and TNFα-treated cells by use of Sytox-green assay which measures the presence of secondary necrotic cells. Consistent with the viability assay results, TNFα, and not TGFβ co-treatment with 5-ITu led to a significant increase in cell death as indicated by enhanced Sytox-green fluorescence (Figure 3B). Interestingly, TGFβ-TNFα co-treatment, an established and more effective EMT stimulus behaved more like TNFα-treated cells in both viability and cytotoxicity assays (Figure 3A & 3B). Moreover, the adenosine kinase inhibitor ABT-702 did not affect cell viability in the conditions tested (Figure 3A & 3B). Similar results were observed in PANC1 cells (Figure 3C). These data indicate that the effect of 5-ITu on TGFβ signaling is not linked to RIPK1-mediated cytotoxicity. Flow-cytometry based cell-cycle analyses revealed an effect of 5-ITu on G2/M arrest (Figure 3D & Supplementary Figure S2), which may explain the reduction in the proportion of viable cells observed in the WST-8 assays. Enhanced cytotoxicity upon TNFα and 5-ITu co-treatment is associated with RIPK1 activation-dependent ripoptosome/necrosome assembly (16). Consistent with this, TNFα/5-ITu-mediated cytotoxic response was sensitive to the RIPK1 inhibitor necroststain-1 (Nec-1) (Figure 3E). However, co-treatment with RIPK1 inhibitors Nec-1 or GSK481 did not have an impact on the effect of 5-ITu on TGFβ-induced EMT as indicated by the expression of the EMT markers *SNAI1* and *CDH2* (Figure 3F). These findings clearly establish that 5-ITu interferes with TGFβ-induced EMT, independent of RIPK1 activation and apoptosis/necroptosis.

**Figure 3.**
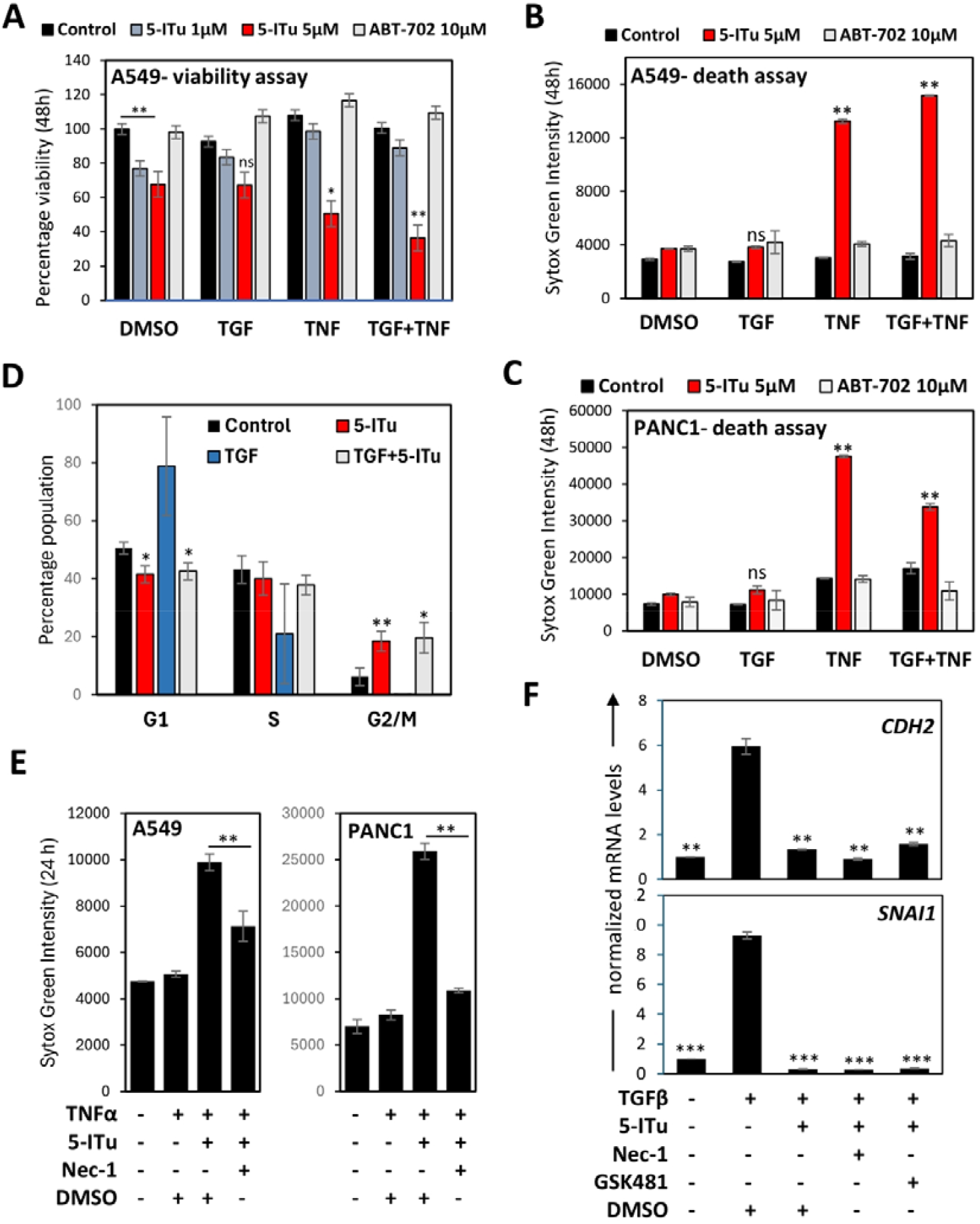
5-ITu mediated suppression of EMT is independent of RIPK1-dependent cytotoxicity. **A**. A549 cells were seeded in a 96 well plate and treated with TGFβ, TNFα in combination with indicated inhibitors or solvent control (DMSO) for 48 h. Cell viability was measured using WST-8. **B-C**. A549 (B) and PANC1 cells (C) were treated with combinations indicated for 48h and cell death was measured and are indicated as Sytox green intensities. **D**. Flow cytometry analysis of cell cycle distribution in A549 cells. Cells were treated as indicated for 24 h, cell cycle phase distribution was calculated and presented (n=3, mean±SD plotted). **E**. A549 (left) and PANC1 cells (right) were treated with combinations indicated for 24 h and cell death was measured and are indicated. F. A549 cells were stimulated with TGFβ in combination with indicated inhibitors or solvent control (DMSO) and EMT marker expression was monitored by qPCR (mean±SD plotted). (* denotes p-value ≤ 0.05, ** denotes p-value ≤ 0.01, *** denotes p-value ≤ 0.001).

### The effect of 5-ITu on EMT is associated with the inhibition of IKK/NFκB signaling

To further delineate the mechanisms of 5-ITu-mediated suppression of EMT, we investigated the NFκB activation status in the cells undergoing TGFβ-induced EMT. Cells treated for 24 h with TGFβ showed significant amounts of IκBα protein and phosphorylated NFκB/p65. However, IκBα and phospho-p65 levels were not significantly affected compared to control cells (Figure 4A). Interestingly, 5-ITu treatment led to a strong suppression of p65 phosphorylation, indicating inhibition of IKK activity and suppression of NFκB-dependent transcription activity. Interestingly, TGFβ receptor-kinase inhibition by SB431542 was associated with enhanced p65 phosphorylation and IκBα-phosphorylation-induced band-shift. We then asked the question whether inhibition of NFκB signaling by IKK1/2 inhibitors will phenocopy the effect of 5-ITu on EMT. The IKK inhibitor BMS345541 suppressed the morphological changes induced by TGFβ in A549 and PANC1 cells (Figure 4B). Consistent with the effect on the EMT-related morphological changes, IKK inhibition also suppressed the EMT-associated gene expression changes at RNA and protein levels (Figure 4C & 4D; Supplementary Figure S3A & S3B), albeit with reduced efficacy compared to 5-ITu. To conclusively prove a role for 5-ITu in the suppression of NFκB-dependent transcription, we performed reporter assays using an NFκB-response element driven reporter construct. As expected, TNFα treatment led to a strong upregulation of NFκB-dependent luciferase activity, while TGFβ induced the reporter activity to a lesser extent (Figure 4E). Consistent with the effects observed on p65 phosphorylation, 5-ITu treatment strongly suppressed TNF and TGFβ-induced reporter activity, similar to the IKKα/β (IKK1/2) inhibitor BMS345541 (22).

**Figure 4.**
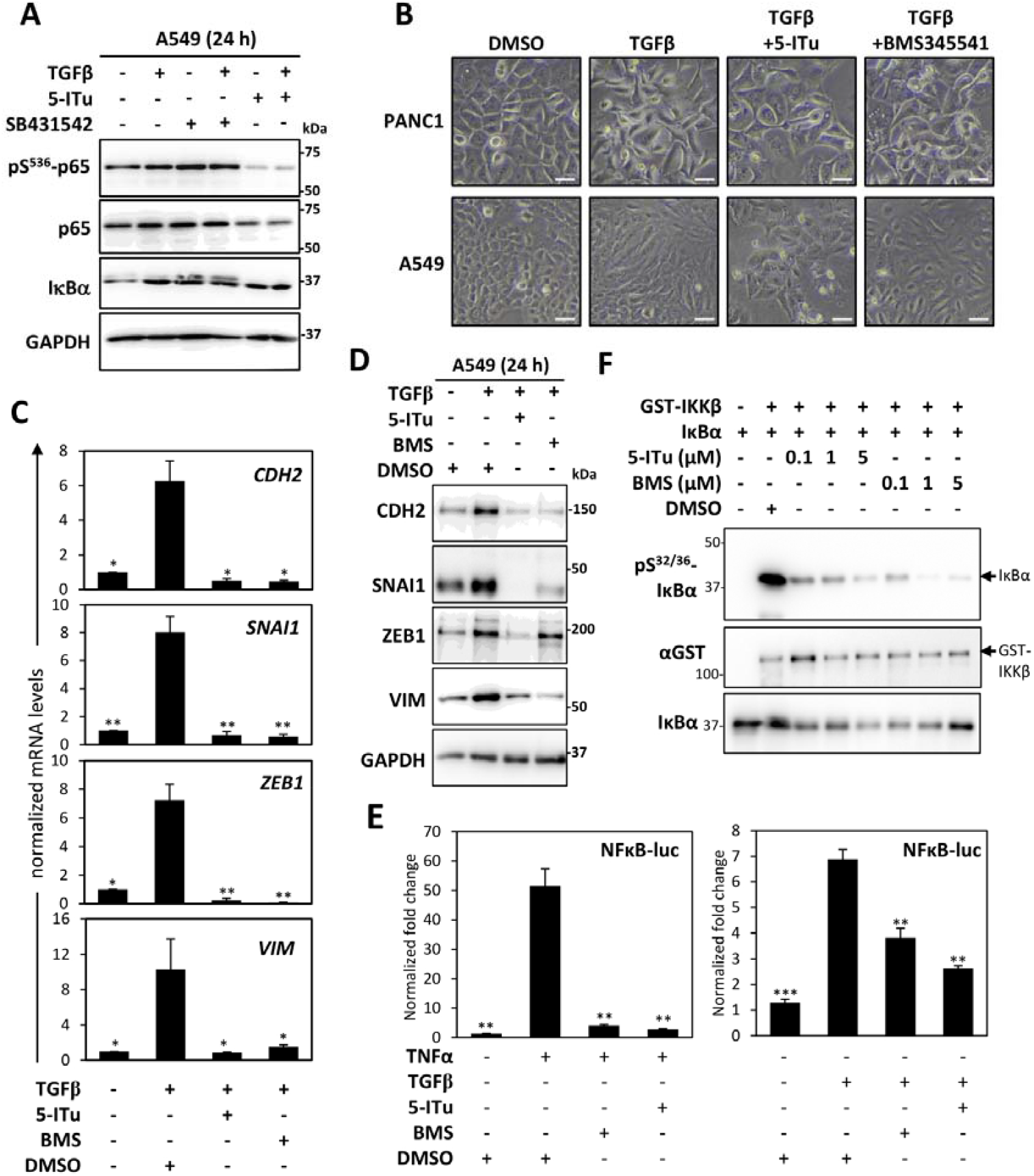
5-ITu inhibits EMT by directly targeting IKKβ kinase activity and NFκB-dependent transcription. **A**. A549 cells were treated with TGFβ in combination with the inhibitors 5-ITu and SB42 for 24h and lysates were probed with pS^536^-p65, total p65 and IκBα to show the effect of 5-ITu on the NFκB pathway. **B**. PANC1 and A549 cells were treated as indicated for 24 h and images were acquired using a phase-contrast 10x objective. Scale bars indicate 50µm. **C-D**. A549 cells were stimulated with TGFβ in combination with indicated inhibitors or solvent control (DMSO) and EMT marker expression was monitored by qPCR (mean±SD plotted) (**C**) and immunoblotting (**D**). **E**. HEK293T cells transfected with luciferase reporter plasmid and Renilla luciferase were treated as indicated for 6h. Luminescence was measured in microplate reader and calculated normalized relative luciferase activity. **F**. *In vitro* kinase assay performed by incubating the kinase IKKβ and the substrate IκBα along with 3 different concentrations of 5-ITu and BMS345541 as indicated for 30 mins. SDS gel was run for the reaction mixes and probed with pS32/S36-IκBα versus total IκBα and with GST. (* denotes p-value ≤ 0.05, ** denotes p-value ≤ 0.01, *** denotes p-value ≤ 0.001).

### IKKβ is a direct protein kinase target of 5-ITu, as demonstrated by *in vivo* kinase assays

Earlier efforts to identify the direct target of 5-ITu relevant to its effect on IKK signaling were unsuccessful (16). Considering that the effect of 5-ITu on IKK/NFκB signaling is conserved downstream to TGFβ as well as TNFα receptors, we hypothesized that 5-ITu could be directly binding IKK and targeting the IKK catalytic activity. To test this hypothesis, we performed *in vitro* kinase assay for IKKβ using recombinant IκBα as substrate. BMS345541, a well-characterized IKKβ inhibitor was used as positive control (22). Interestingly, 5-ITu dose-dependently inhibited IκBα phosphorylation with an efficiency comparable to that of BMS345541 (Figure 4F; Supplementary Figure S3C). These findings indicate that 5-ITu-mediated inhibition of TGFβ-mediated EMT is a consequence of direct IKK inhibition by 5-ITu, resulting in the suppression of NFκB signaling.

### A 5-ITu sensitive NFκB-dependent gene signature relevant to EMT and cancer

TGFβ-mediated EMT is mainly regulated by the SMAD2/3-dependent signaling pathways altering the expression of EMT-related transcription factors and cadherin family proteins. Previous studies have established a SMAD-independent role for NFκB signaling in the regulation of EMT (5, 34). This is consistent with the effect of IKK inhibitor BMS345541 on TGFβ-induced EMT and expression of EMT-related genes (Figure 4B-D). To further confirm that the effect of 5-ITu on EMT is mediated by the suppression of NFκB-dependent gene transcription, we compared the gene expression signatures unique to TGFβ signaling and that of TNF-NFκB pathway. The gene sets for TGFβ and TNFα/NFκB pathways were obtained from the Molecular Signatures Database (www.gsea-msigdb.org)(28, 29) and consisted of 54 and 200 genes respectively. There were ten genes common between the two datasets indicating these are potential NFκB-dependent transcripts downstream to TGFβ signaling (Figure 5A). Interestingly, expression of this 10 gene signature was associated with overall survival in lung cancer (Figure 5B). Similar effects were also observed in case of pancreatic cancer (Supplementary Figure S4A). We then asked the question whether these genes are sensitive to 5-ITu treatment. While four of the genes (*SMAD3, SERPINE1, TGIF* and *PMEPA1*) displayed TGFβ-induced upregulation as well as suppression by 5-ITu treatment (Figure 5C), three genes (*BMP2, KLF10* and *IFNGR2*) were not expressed to detectable levels in A549 cells, independent of TGFβ stimulation. While *JUNB* displayed a reverse tendency, the expression changes were not statistically significant, and two genes (*ID2* and *PPP1R15A*) did not display significant alterations in response to TGFβ or 5-ITu treatment (Figure 5D). We also monitored the effect of 5-ITu on the expression of *XIAP*, a target and regulator of NFκB pathway as well as EMT (35). Moreover, 5-ITu treatment also inhibited TGFβ-induced *XIAP* expression (Figure 5E). Interestingly, the gene signature consisting of the four 5-ITu sensitive genes was enough to predict overall survival in lung (Figure 5F) and pancreatic cancer (Supplementary Figure S4B). Moreover, higher expression of these four genes was associated with poorer disease-free survival (Figure 5G & H), revealing the prognostic significance of the 5-ITu sensitive gene signature.

**Figure 5.**
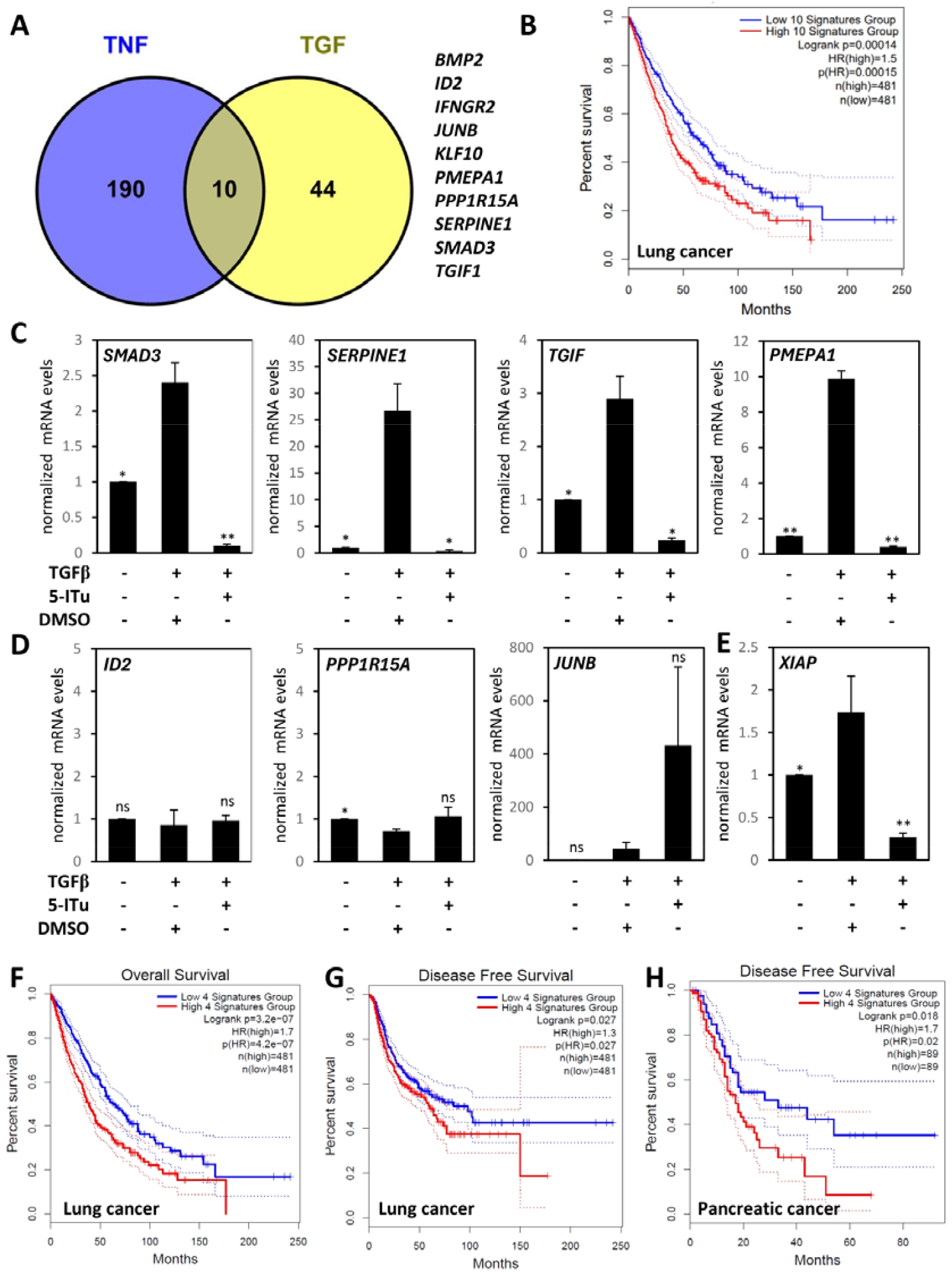
A NFκB-dependent, 5-ITu-sensitive gene signature predicts disease-free survival in cancer. **A**. A Venn diagram depicting the common genes in the hallmark genes of TNF/NFκB and TGFβ signaling. The 10 common genes are shown in right panel. **B**. The 10-gene signature predicts overall survival in lung cancer. **C-E**. A549 cells were treated with TGFβ alone or in combination with 5-ITu for 24h and expression of the indicated transcripts were analyzed by qRT-PCR (* denotes p-value ≤ 0.05, ** denotes p-value ≤ 0.01, *** denotes p-value ≤ 0.001). **F**. Kaplan-Meier plot shows the prediction of overall survival in lung cancer by the 5-ITu-sensitive 4-gene signature. **G & H**. Kaplan-Meier plots show the prediction of disease-free survival in lung cancer (G) and pancreatic cancer (H) by the 5-ITu-sensitive 4-gene signature.

## Discussion

We had earlier identified 5-ITu in a screen for small molecule inhibitors which sensitizes MK2-KO MEFs to TNF-induced necroptosis (16). 5-ITu was shown to facilitate RIPK1 activation by compromising an IKK1/2-mediated checkpoint on RIPK1 activation. Here we present evidence for 5-ITu as a strong inhibitor of type-II EMT induced by TGFβ. Consistent with the effect on TNFα and TLR4 signaling, the effect of 5-ITu on IKK signaling was also conserved in case of TGFβ-induced EMT. Unlike in the case of TNFα/LPS, 5-ITu did not enhance cytotoxicity in combination with TGFβ. However, 5-ITu treatment significantly affected viable cell numbers and cell cycle progression independent of TGFβ. This is in line with the role of 5-ITu as a nucleoside analog and potential chemotherapeutic agent. As established in the literature, TGFβ is a weak activator of NFκB signaling compared to TNFα and co-treatment with TNFα enhances TGFβ-induced EMT, by activating strong NFκB signaling (36). This is evident from the NFκB-luciferase reporter assays which showed almost 10-fold difference in the induction of reporter activity between TNFα and TGFβ (Figure 4E). Despite weak activation of NFκB pathway by TGFβ, NFκB-dependent transcription is key to the EMT program. Consistent with this, IKK2 inhibition by specific inhibitor BMS345541 suppressed EMT like 5-ITu, and 5-ITu treatment downregulated NFκB/p65 phosphorylation concomitant with inhibition of TGFβ-induced EMT. Interestingly, in immunoblots of samples from cells treated with TGFβ-receptor kinase inhibitor SB431542, we observed the accumulation of a shifted band for IκBα and enhanced phospho-p65 levels (37). Band-shifts on SDS-PAGE gels are often associated with protein phosphorylation in basophilic motifs and similar shifts of IκBα has been attributed to S32/S36 phosphorylation (38). Such crosstalk between long-term inactivation of TGFβ receptor activity and hyperactivation of IKK signaling has not been reported previously and warrants further investigation.

EMT is essential for embryonic development and is involved in cancer progression and metastasis (39, 40). EMT-associated transcription factors like SNAI1, SNAI2 and ZEB1 are implicated in several forms of cancer (41-44). While the role of NFκB signaling in cancer has been investigated more in the context of inflammation induced carcinogenesis, survival signaling and suppression of apoptotic pathways (45), NFκB-dependent transcription also play key roles in EMT, metastasis and development of more aggressive and invasive cancers (46). Interestingly, a recent study used the presence of high NFκB activity in the cancer cells as a strategy for activating NFκB-dependent CRISPR/Cas13a in colon tumors to suppress oncogenes by therapeutic AAV (adeno-associated virus) vectors (47). PD-L1 expression in cancer cells and tumor associated myeloid cells promote tumor cell survival by preventing lymphocyte activation (48). PD-L1 is hence a major target for immunotherapy using immune-checkpoint inhibitors. Interestingly, PD-L1 expression is NFκB-dependent, is upregulated during EMT in lung cancer cells and over-expression of PD-L1 in tumors is often associated with the upregulation of EMT markers (49, 50)

TNFα/NFκB is an important SMAD-independent signaling pathway which modulates cancer cell EMT, driven by the cocktail of secreted factors released by cancer cells and tumor-associated macrophages (51). Unlike in TNFα-dependent cell death, we do not see the involvement of RIPK1 activation in 5-ITu-mediated suppression of TGFβ-induced EMT. While the sensitizing effect of 5-ITu on necroptosis is linked to the inactivation of IKK-dependent checkpoint on RIPK1 activation, the effect on EMT is independent of RIPK1 and potentially linked to NFκB-dependent transcription of EMT genes like *SNAI1*. Interestingly, we identified a 4-gene signature defined by (i) TNF/NFκB-dependent expression, (ii) regulation by TGFβ signaling and (iii) sensitivity to 5-ITu which is effective in predicting disease-free survival in lung and pancreatic cancer. A previous study had reported a larger gene expression signature for prostate cancer prognosis, which consisted of 21 NFκB-dependent genes (51). Another gene signature of prognostic value identified for low-grade gliomas also consisted of 14 NFκB-dependent genes (52). We have focused on lung/pancreatic cancer, as our cell culture data is derived from cancer cells of pancreatic and lung cancer origin. However, this approach and the 4-gene signature identified in our study could be applicable to other forms of cancer.

5-ITu was discovered as an adenosine kinase inhibitor and was later established as a chemotherapeutic molecule with antineoplastic properties in mouse model *in vivo* (6, 7). While some effects of 5-ITu, including the blockade of SARS-CoV2 replication is associated with its ability to inhibit adenosine kinase activity, other effects are independent of adenosine kinase inhibition (8). Later studies established ERK2, Haspin and several others as protein kinase targets of 5-ITu (10, 13). The role of 5-ITu in TNFα and TGFβ signaling is indeed independent of adenosine kinase inhibition. Our earlier efforts to identify the direct kinase target of 5-ITu, relevant to its effect on TNF-induced cytotoxicity was unsuccessful, despite identifying a panel of potential 5-ITu targets (16). Using *in vitro* kinase assays, we have now identified IKK2/IKKβ as a direct protein kinase target of 5-ITu. These results indicate that the effect of 5-ITu on TNFα-induced RIPK1 activation and TGFβ-induced EMT are mediated by direct inhibition of the IKK1/2 activity. Incidentally, IKKs were not included in the earlier screens for 5-ITu targets (13, 16). While several other targets of 5-ITu, including protein kinases are relevant to chemotherapeutic intervention against cancer, the sensitization of cancer cells to TNF/smac-mimetics induced necroptosis and interference with EMT are interesting candidate features for anticancer drugs. Further *in vivo* studies probing the effect of 5-ITu alone and in combination with smac-mimetics will be necessary to delineate the efficacy and mechanistic contribution of RIPK1-dependent necroptosis and EMT inhibition on the anti-tumor activity of 5-ITu and/or analogous compounds.

## Supporting information

Supplementary Material-Uncropped Immunoblots

Supplementary Table 2 - Statistical Source data

## Acknowledgements

S.D. thanks Department of Biotechnology (DBT), India for Ramalingaswami Re-entry Fellowship (BT/ RLF/Re-entry/10/2019) and DBT Wellcome Trust India Alliance for Intermediate Fellowship support (IA/I/22/2/506497). V.G. was supported by the DBT Wellcome Trust India Alliance grant. M.B.M. thanks IIT Delhi for intramural grant support (MFIRP/MI02714). S.B. thanks Department of Education, Government of India for PhD fellowship. M.M. acknowledges support from intramural funds of CSIR-IGIB (OLP2303). The authors thank Kusuma School of Biological Sciences and the Central Research Facility (CRF), IIT Delhi for infrastructural support, infrared imager facility and confocal microscopy facility. We thank Dr. Shantanu Chowdhury lab and CSIR-IGIB for infrastructural support. The authors thank Dr. Mark Windheim (MHH, Hannover) for the gift of NFκB-luciferase reporter plasmid. Authors thank the TCGA Research Network and the GEPIA2 server for access to the tumor gene expression datasets.

## Author Contribution Statement

S.B. performed majority of the experiments, designed experiments, and analysed data. M.M. performed EMT marker analyses. V.G. performed the cell cycle experiments and tested the effect of RIPK1 inhibitors on EMT markers. S.D. & M.B.M. procured funding, designed and supervised experiments and analysed data. S.D., S.B. and M.B.M. prepared the manuscript.

## Data Availability Statement

All data generated during this study leading to the findings presented here are included in this published article and its supplementary data files. The patient survival data from The Cancer Genome Atlas (TCGA) was accessed, and survival plots were generated directly using the GEPIA portal.

## Ethics Approval

This study doesn’t involve use of animals or human subjects and are exempt from ethical approvals. The patient dataset analysed were publicly available from the TCGA portal.

Consent to participate Not applicable

