## Supplementary figures and images for "5-Iodotubercidin inhibits Epithelial to Mesenchymal Transition by inhibiting IKK/NFκB-dependent gene expression"

### Supplementary Material-Uncropped Immunoblots

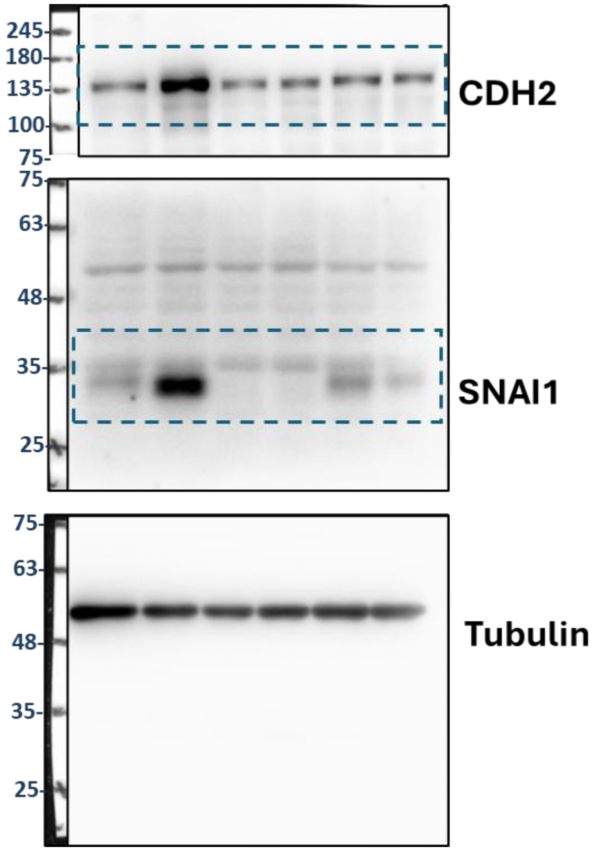

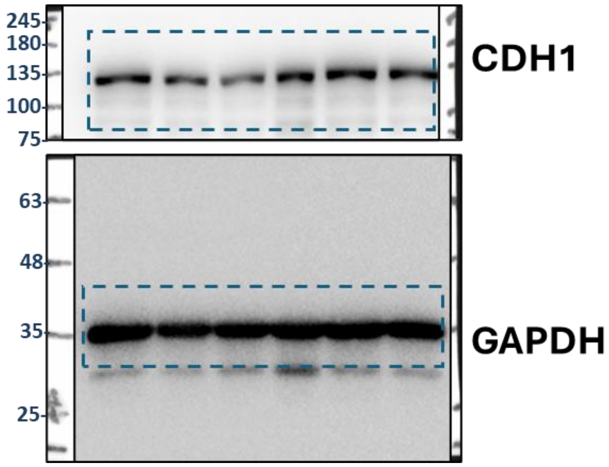

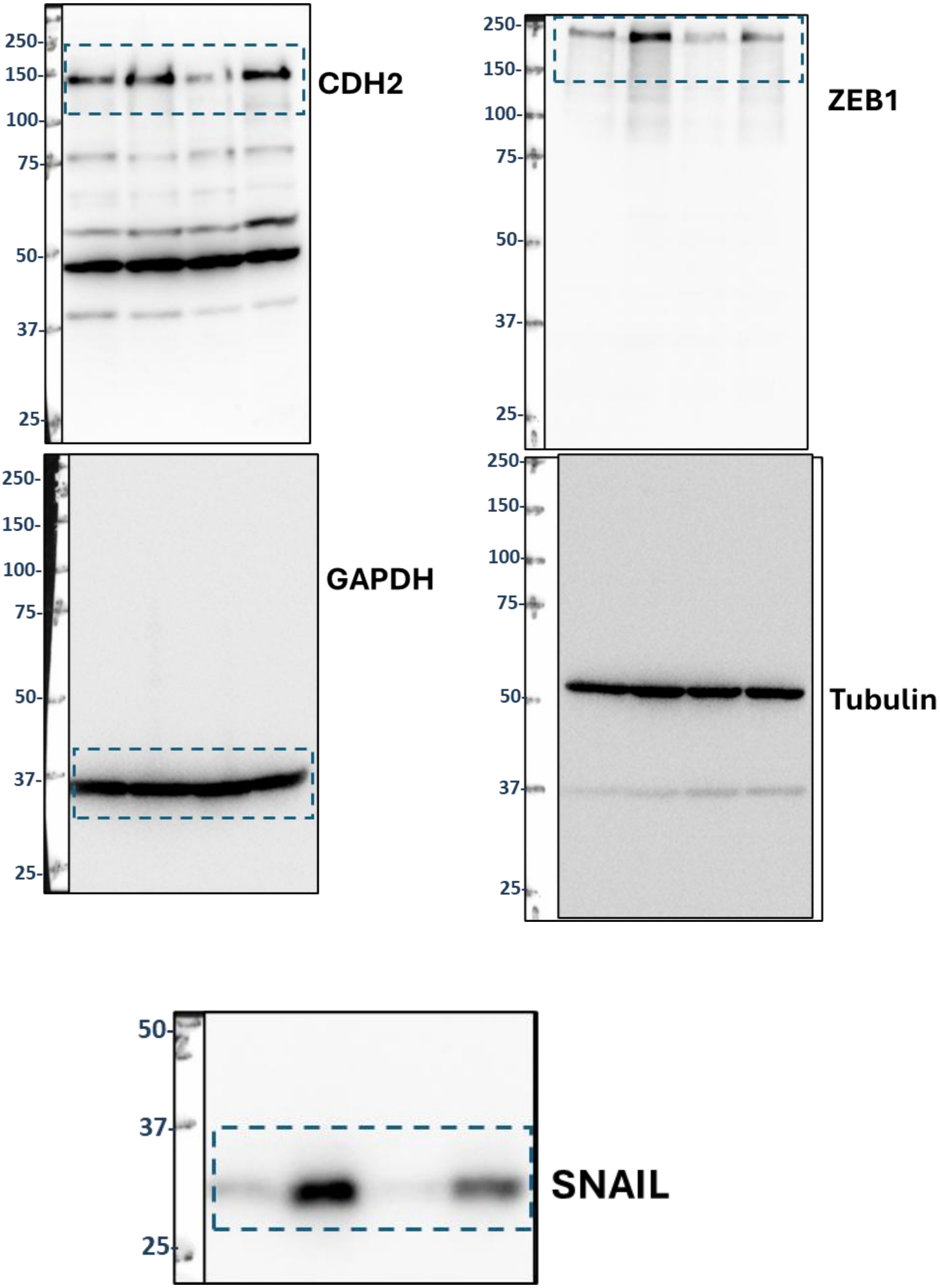

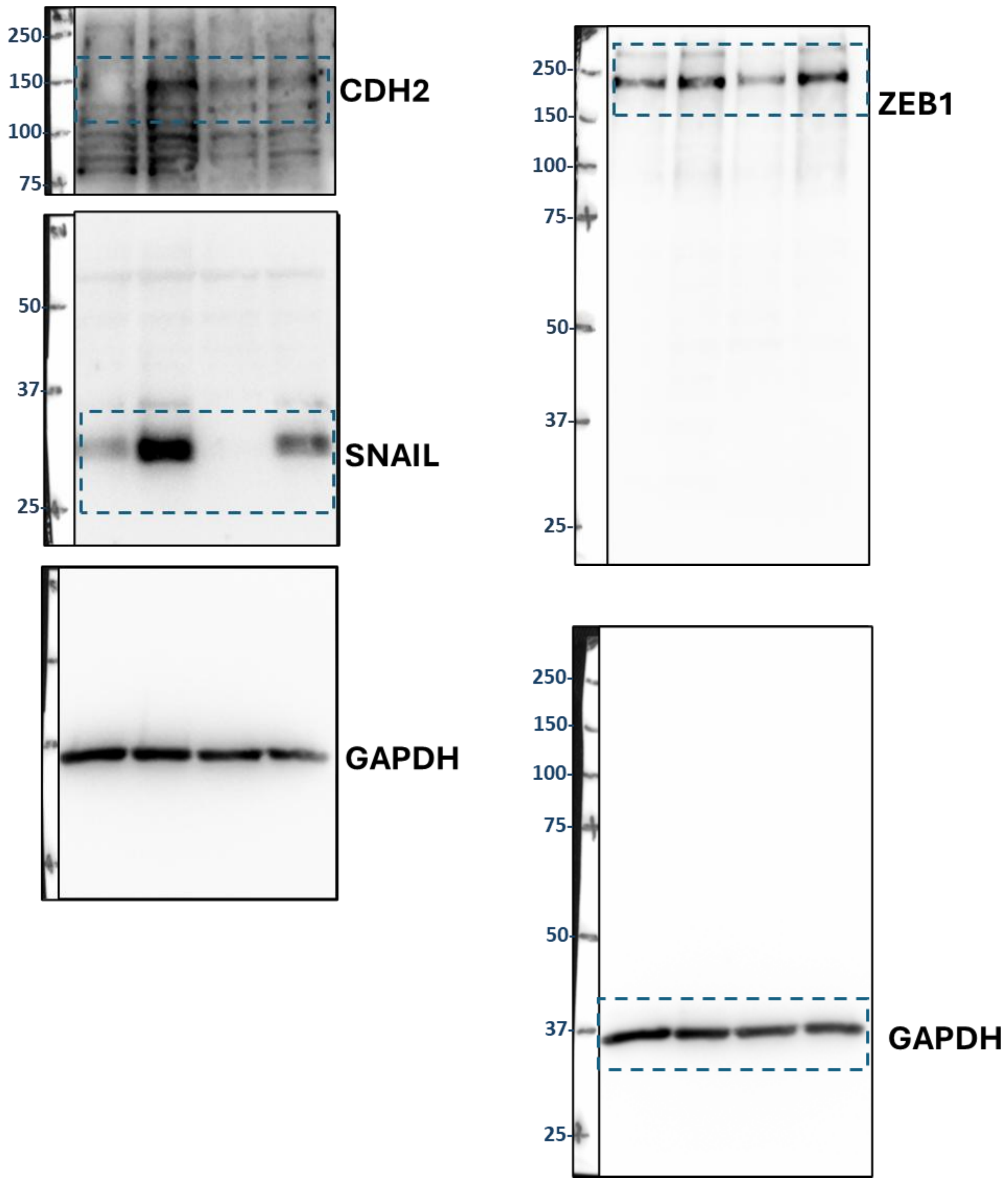

#### Figure 4A

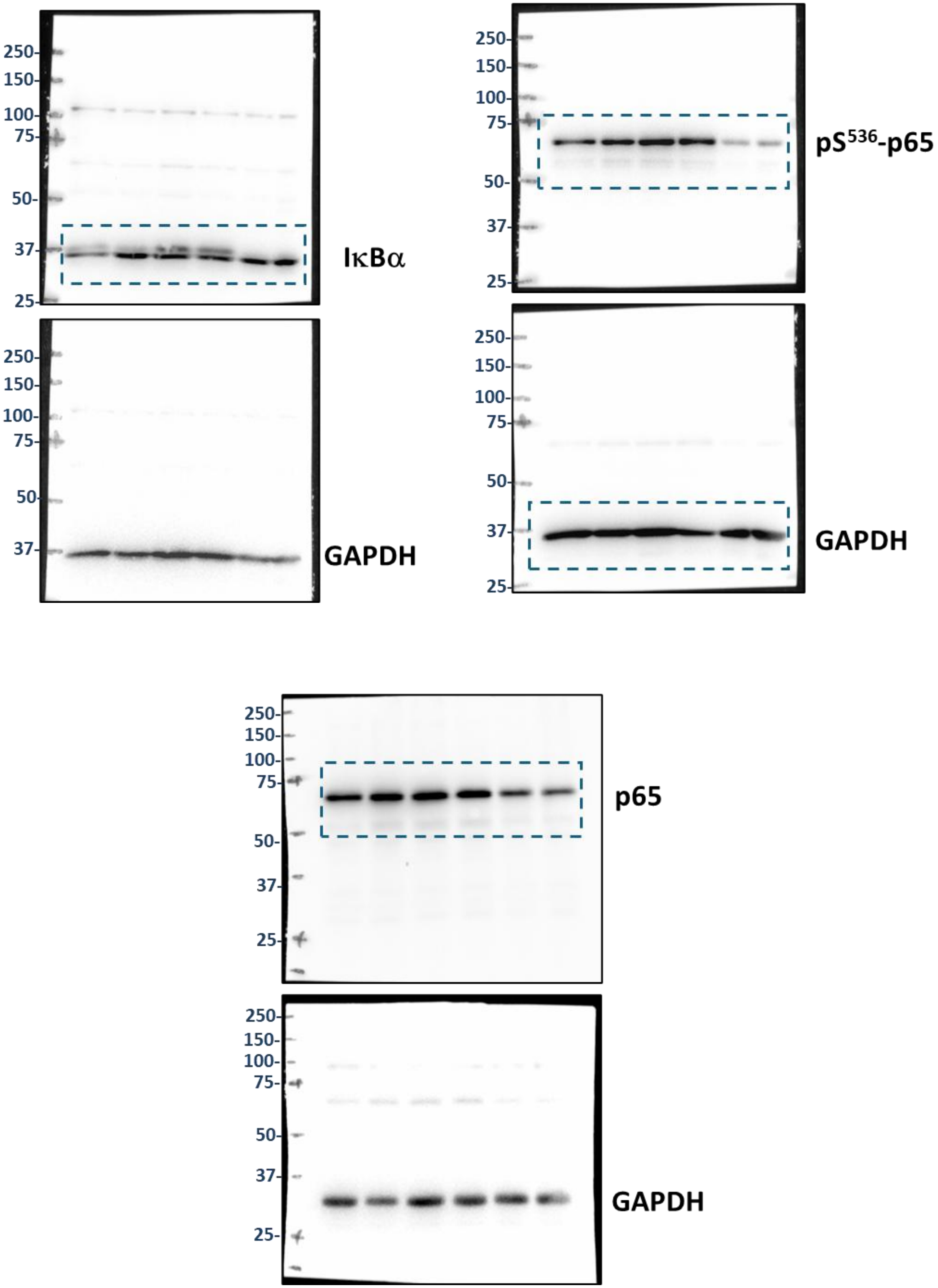

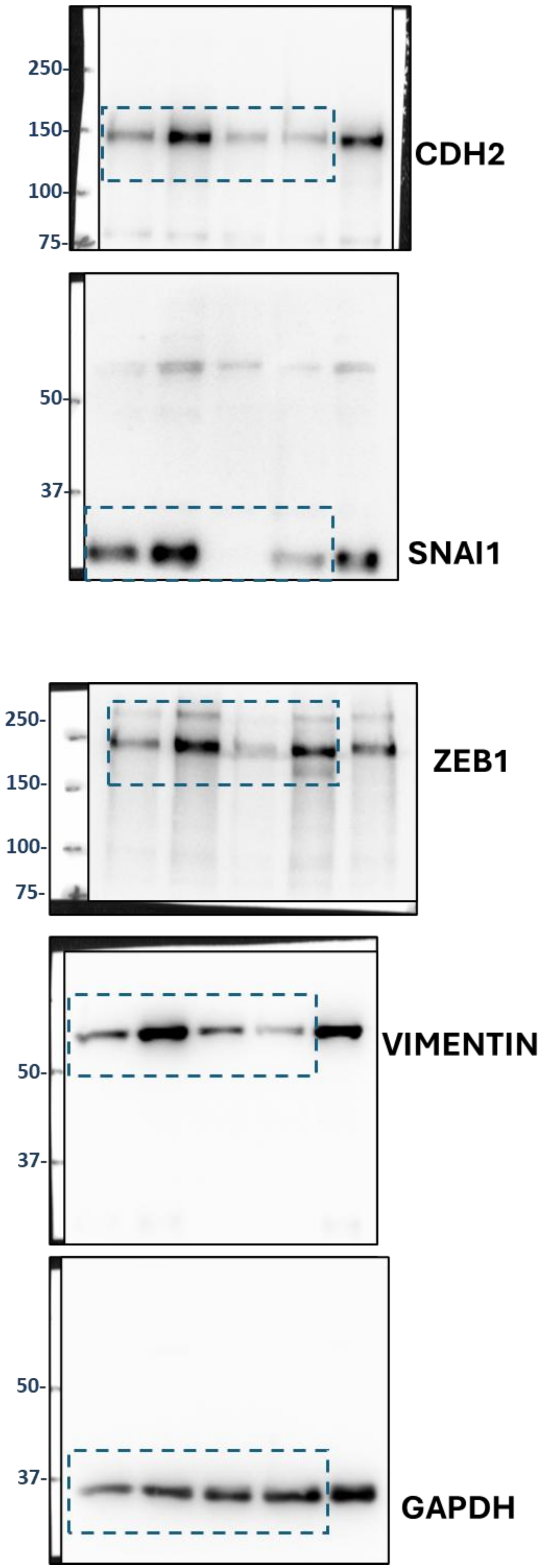

**Figure 4F**

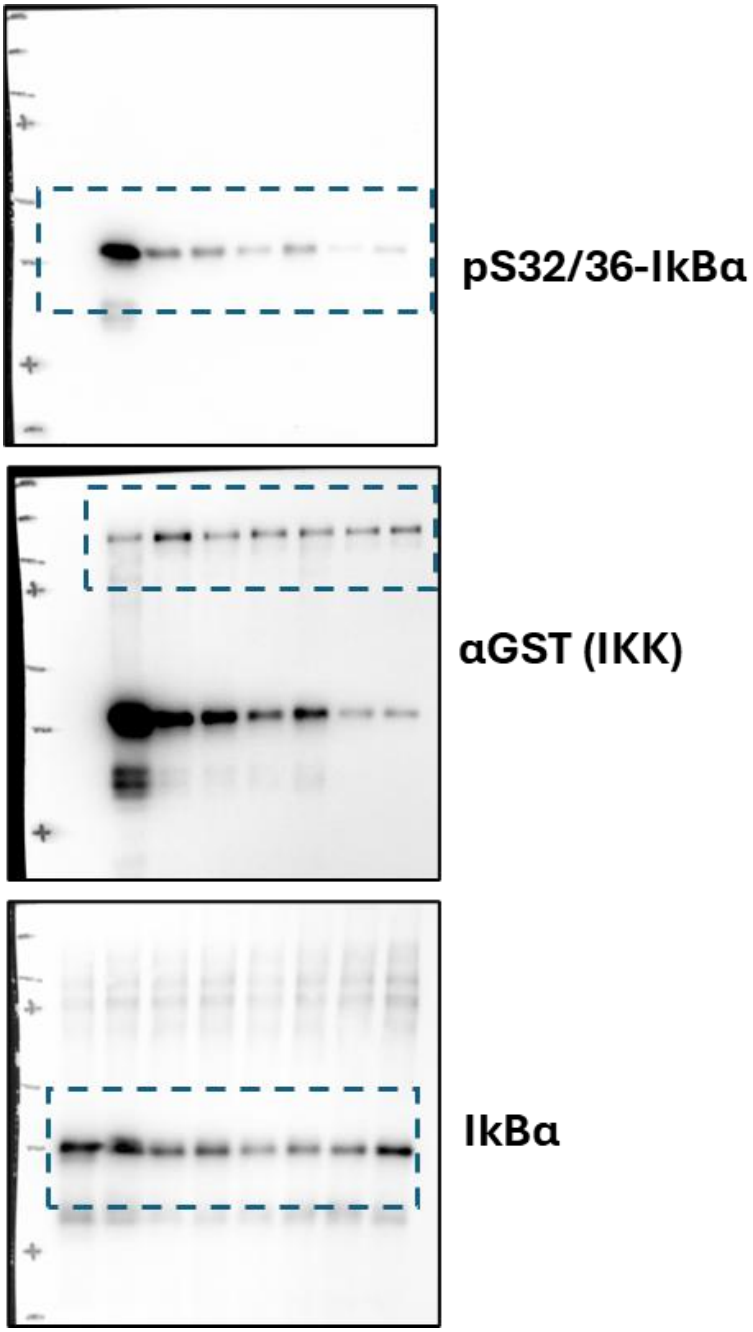

##### Suppl. Figure S3B

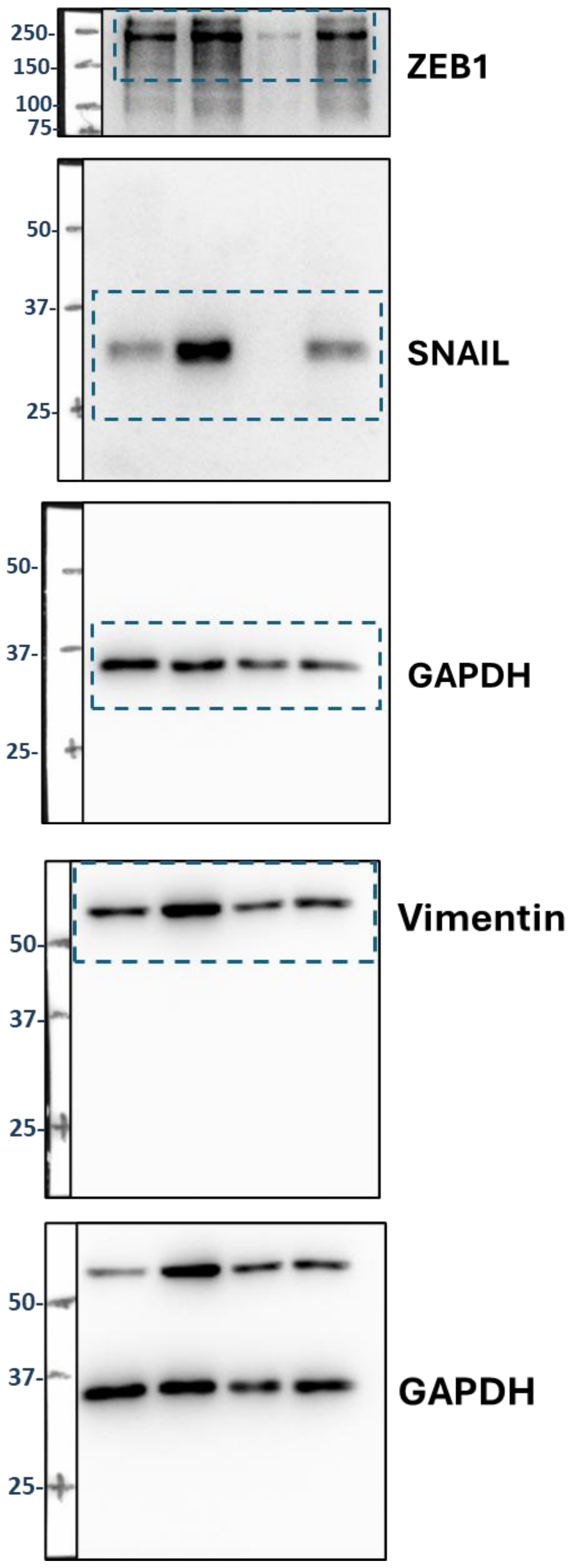
