## Supplementary Figures S1 - S4 for "5-Iodotubercidin inhibits Epithelial to Mesenchymal Transition by inhibiting IKK/NFκB-dependent gene expression"

### Supplementary Figure S1

**A**

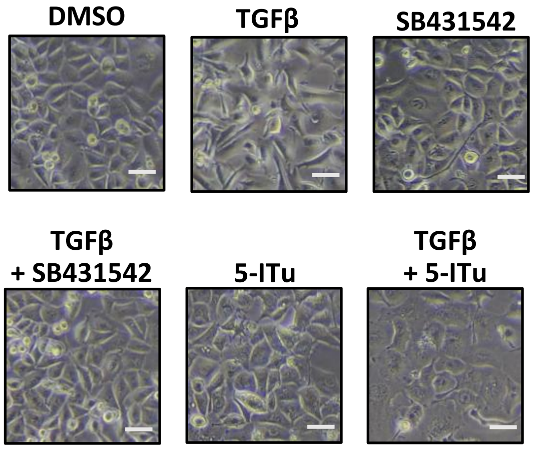


**B**

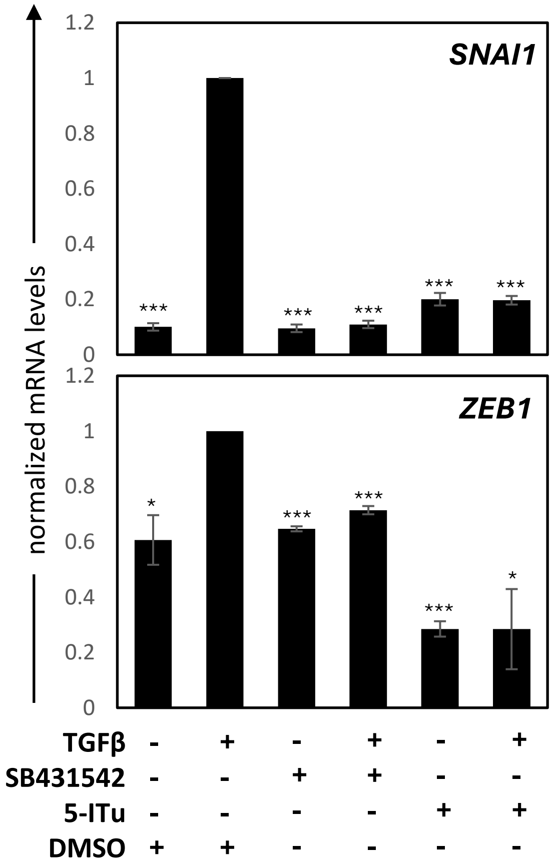

**Supplementary Figure S2**

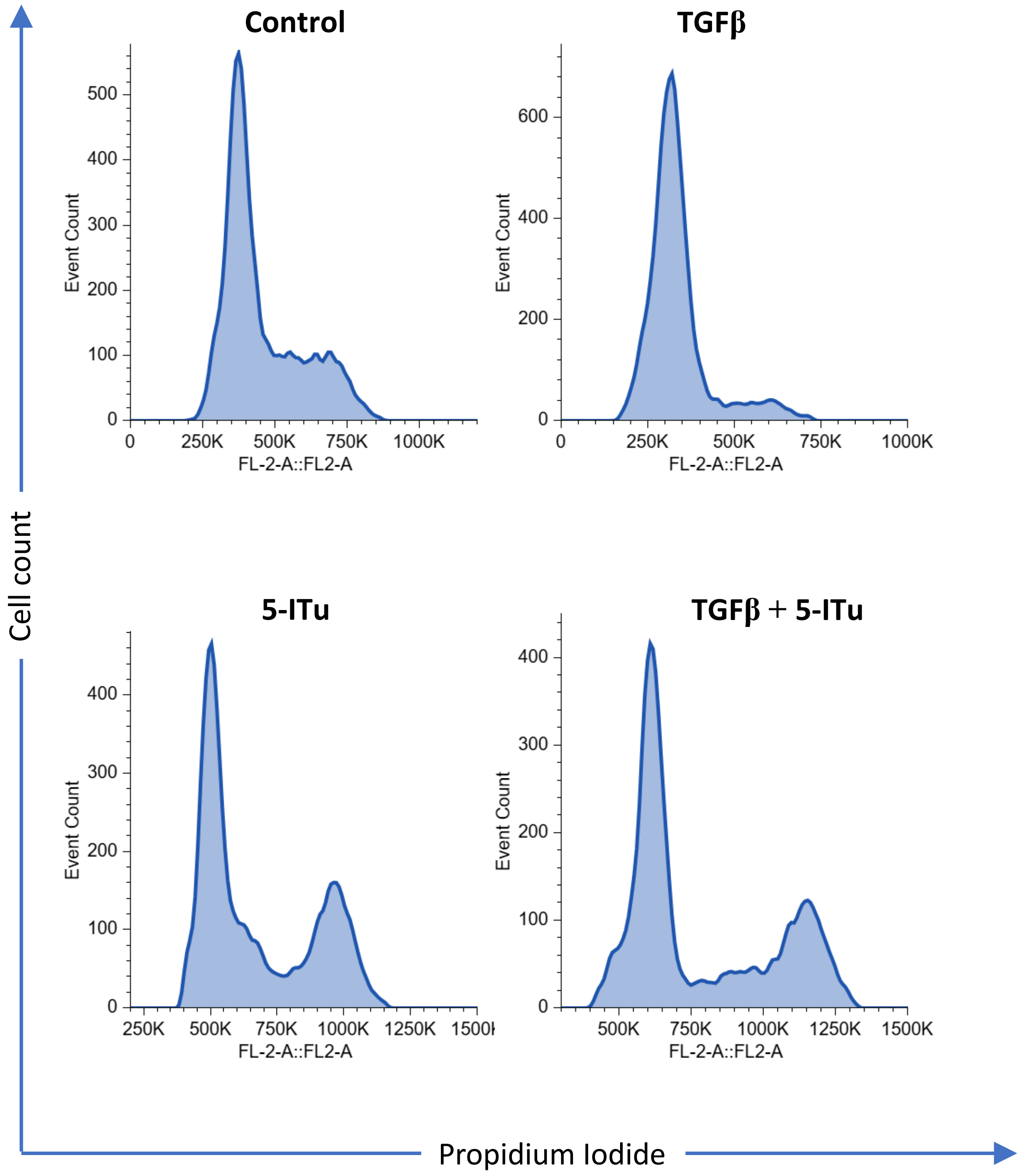

### Supplementary Figure S3

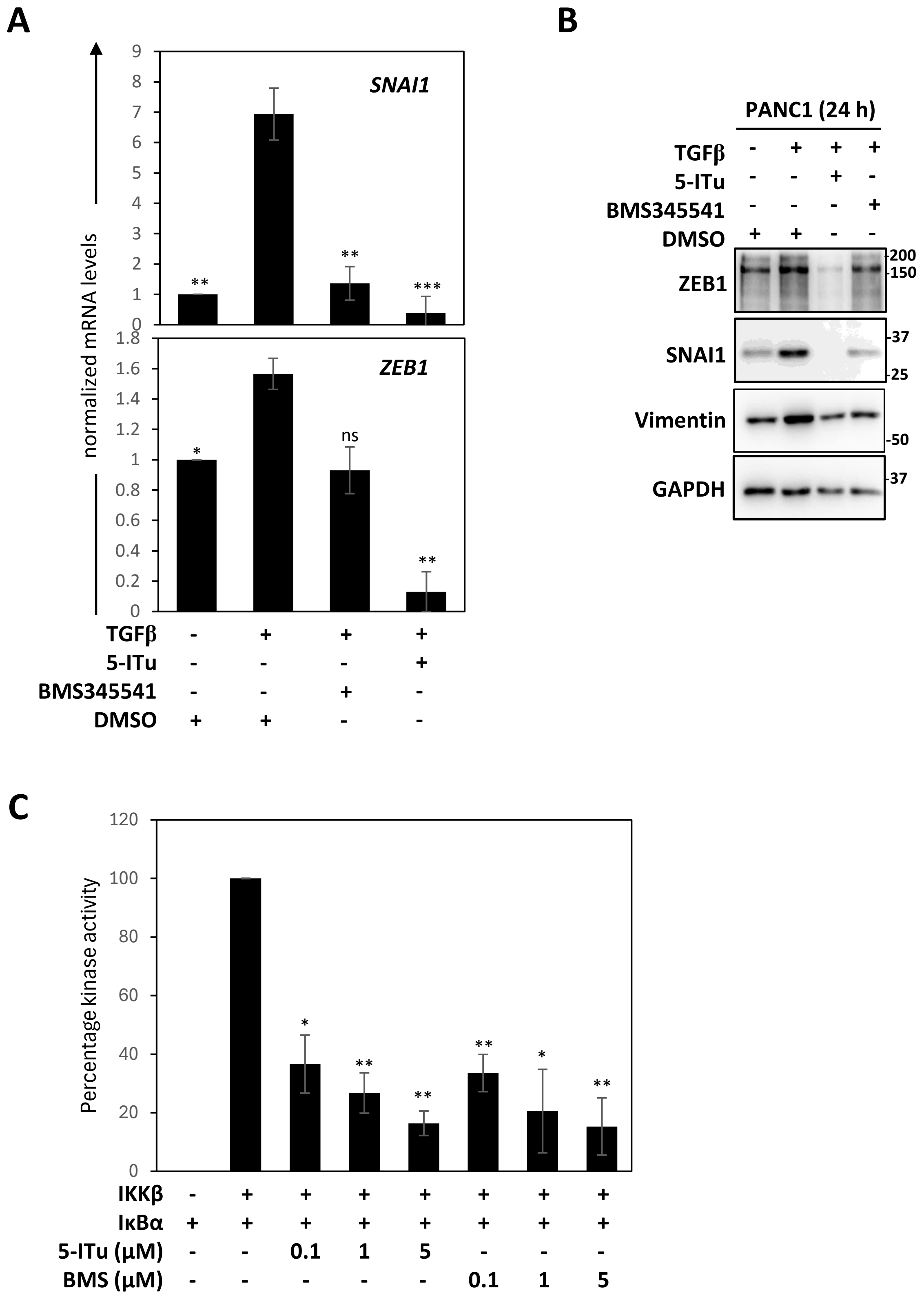

### Supplementary Figure S4

A

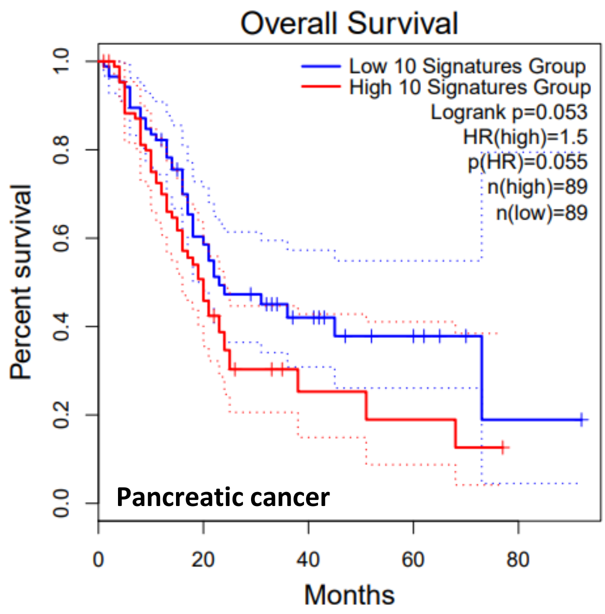


B

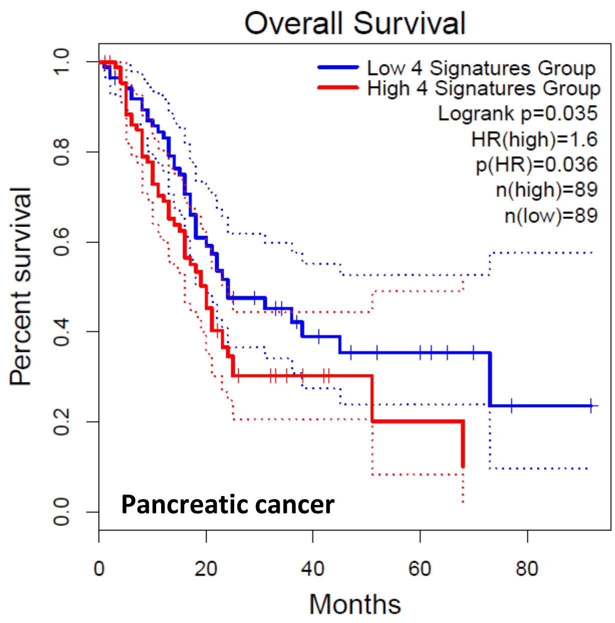
