## Supplementary Files Information for "5-Iodotubercidin inhibits Epithelial to Mesenchymal Transition by inhibiting IKK/NFκB-dependent gene expression"

### Supplementary Information Legends

1. **Supplementary Figures S1-S4 (single .pdf file)**

**Supplementary Figure 1. 5-ITu inhibits TGFβ-induced EMT in PANC1 cells. A.** PANC1 cells were treated as indicated for 24 h and images were acquired using a phase-contrast 10x objective. Scale bars indicate 50 µm. **B.** PANC1 cells were stimulated with TGFβ in combination with indicated inhibitors or solvent control (DMSO) and EMT marker (*ZEB1, SNAI1*) expression was monitored by qPCR (n=3, mean±SD for 3 independent experiments) (* denotes p-value ≤ 0.05 and *** denotes p≤ 0.001).

**Supplementary Figure 2. Cell-cycle analyses of 5-ITu-treated A549 cells.** Representative flow-cytometry histograms for the data presented in Figure 3D. Cells were treated for 24h as follows: untreated control cells, cells treated with 5 µM 5-ITu, 5 ng/mL TGF-β or both, and fixed and stained with propidium iodide. DNA content was analyzed by flow cytometry.

**Supplementary Figure 3. IKK inhibition and Epithelial-Mesenchymal Transition. A-B.** PANC1 cells were treated as indicated and EMT marker expression was monitored using qRT-PCR (A) and immunoblotting (B). **C.** Quantitative data of IKK2 kinase assays represented in Figure 4F. Data is represented as percentage activity, value quantified as $pS^{32/S36}$- IκBα band intensities normalized to total IκBα. (n=3, mean±SD for 3 independent experiments) (* denotes p-value ≤ 0.05, ** denotes p-value ≤ 0.01 and *** denotes p-value ≤ 0.001).

**Supplementary Figure 4. NFκB/5-ITu-dependent prognostic gene expression signatures in pancreatic cancer. A & B.** TCGA pancreatic datasets were used to plot overall survival of pancreatic cancer patients based on the 10-gene signature mentioned in Figure 5A-B (A) or the NFκB-dependent, 5-ITu-sensitive 4-gene signature (B).

2. **Supplementary Table S1. Sequence of primers used in the study.** Sequence details of qPCR primers used in the study (PDF file).

3. **Supplementary Table S2. Statistics and source data for the quantitative results.** The actual values (source data) and the *p*-values calculated for the quantitative data presented in the manuscript (PDF file)

4. **Supplementary Data File 2: Uncropped blots** (PDF file)
