## Supplementary Table 2 - Statistical Source data for "5-Iodotubercidin inhibits Epithelial to Mesenchymal Transition by inhibiting IKK/NFκB-dependent gene expression"

#### Figure 1C

0.05*, P < 0.01**, and P < 0.001***

| CDH1 | REP 1 | REP 2 | REP 3 | | AVERAGE | ST DEV | p Value | |
|---|---|---|---|---|---|---|---|---|
| CTRL | 1 | 1 | 1 | | 1 | 0 | 0.000577 | *** |
| TGF β | 0.418028 | 0.452734 | 0.406919 | | 0.425893 | 0.019513 | | |
| SB42 | 0.908241 | 1.19469 | 1.149993 | | 1.084308 | 0.125828 | 0.015975 | * |
| SB42+TGFβ | 0.928137 | 1.2107 | 1.173223 | | 1.10402 | 0.125306 | 0.014903 | * |
| 5-ITu | 0.65516 | 0.812188 | 0.941464 | | 0.802937 | 0.117066 | 0.041707 | * |
| 5-ITu+TGF β | 0.661144 | 0.703288 | 0.863423 | | 0.742619 | 0.087137 | 0.030761 | * |

| CDH2 | REP 1 | REP 2 | REP 3 | | AVG | ST DEV | p Value | |
|---|---|---|---|---|---|---|---|---|
| CTRL | 1 | 1 | 1 | | 1 | 0 | 0.003268 | ** |
| TGF β | 5.306714 | 6.203312 | 6.062269 | | 5.857431 | 0.482121 | | |
| SB42 | 0.757755 | 1.03765 | 1.240437 | | 1.011948 | 0.242365 | 0.000633 | *** |
| SB42+TGFβ | 0.867886 | 0.937138 | 1.390887 | | 1.065304 | 0.284081 | 0.000435 | *** |
| 5-ITu | 0.657104 | 0.821462 | 1.180548 | | 0.886371 | 0.267691 | 0.000453 | *** |
| 5-ITu+TGF β | 0.671603 | 0.68561 | 0.751148 | | 0.702787 | 0.042463 | 0.002728 | ** |

| SNAIL | REP 1 | REP 2 | REP 3 | | AVG | ST DEV | p Value | |
|---|---|---|---|---|---|---|---|---|
| CTRL | 1 | 1 | 1 | | 1 | 0 | 0.004941 | ** |
| TGF β | 5.220454 | 4.310947 | 4.956981 | | 4.829461 | 0.467971 | | |
| SB42 | 0.568359 | 1.492136 | 0.915127 | | 0.991874 | 0.466646 | 0.00055 | *** |
| SB42+TGFβ | 0.794211 | 0.571283 | 0.183648 | | 0.516381 | 0.308962 | 0.000424 | *** |
| 5-ITu | 0.106381 | 0.248266 | 0.147125 | | 0.167258 | 0.073054 | 0.00277 | ** |
| 5-ITu+TGF β | 0.080814 | 0.18247 | 0.15286 | | 0.138715 | 0.052283 | 0.002996 | ** |

| ZEB 1 | REP 1 | REP 2 | REP 3 | | AVG | ST DEV | p Value | |
|---|---|---|---|---|---|---|---|---|
| CTRL | 1 | 1 | 1 | | 1 | 0 | 0.025764 | * |
| TGF β | 1.75446 | 1.751204 | 2.194879 | | 1.900181 | 0.255221 | | |
| SB42 | 0.957668 | 1.535287 | 1.358587 | | 1.283847 | 0.295974 | 0.053597 | * |
| SB42+TGFβ | 1.005697 | 1.129339 | 0.589968 | | 0.908335 | 0.282559 | 0.010981 | * |
| 5-ITu | 0.202396 | 0.21325 | 0.351998 | | 0.255881 | 0.083417 | 0.004349 | ** |
| 5-ITu+TGF β | 0.191505 | 0.21325 | 0.180423 | | 0.195059 | 0.016699 | 0.007191 | ** |

#### Figure 1E

**A549**

| Samples | Migration index 18h | Average 18h | Stdev | p value | |
|---|---|---|---|---|---|
| Control | 1.246753 | 1.143679 | 0.097586 | 7.01E-05 | *** |
| | 1.185013 | | | | |
| | 0.985479 | | | | |
| | 1.0793 | | | | |
| | 1.221847 | | | | |
| TGF | 1.586458 | 2.02685 | 0.218269 | | |
| | 1.934804 | | | | |
| | 2.101021 | | | | |
| | 2.212033 | | | | |
| | 2.219735 | | | | |
| | 2.107047 | | | | |
| TGF+SB42 | 0.948109 | 1.232436 | 0.197445 | 0.001233 | ** |
| | 1.470984 | | | | |
| | 1.162743 | | | | |
| | 1.347908 | | | | |
| TGF+5ITu 0.1μM | 1.619126 | 1.835967 | 0.11568 | 0.13144 | ns |
| | 1.884303 | | | | |
| | 1.818943 | | | | |
| | 1.93385 | | | | |
| | 1.923612 | | | | |
| TGF+5ITu 1μM | 1.642807 | 1.615854 | 0.16044 | 0.633284 | ns |
| | 1.665879 | | | | |
| | 1.927248 | | | | |
| | 1.434937 | | | | |
| | 1.512388 | | | | |
| | 1.511862 | | | | |
| TGF+5ITu 5μM | 0.827559 | 0.729856 | 0.106648 | 4.11E-06 | *** |
| | 0.546852 | | | | |
| | 0.671957 | | | | |
| | 0.790803 | | | | |

| | | | | | |
|---|---|---|---|---|---|
| | 0.81211 | | | | |
| TGF+5ITu 10µM | 0.756752 | 0.524393 | 0.242381 | 1.32E-06 | *** |
| | 0.891019 | | | | |
| | 0.438893 | | | | |
| | 0.397705 | | | | |
| | 0.149445 | | | | |
| | 0.512546 | | | | |

**PANC1**

| Samples | Migration index 24 h | Average | Stdev | p value | |
|---|---|---|---|---|---|
| Control | 1.563389 | 1.563389 | 0.153712 | 6.97E-05 | *** |
| | 1.325607 | | | | |
| | 1.10615 | | | | |
| | 1.212199 | | | | |
| | 1.361349 | | | | |
| TGF | 1.848338 | 2.36608 | 0.2695 | | |
| | 2.432182 | | | | |
| | 2.232079 | | | | |
| | 2.533784 | | | | |
| | 2.6974 | | | | |
| | 2.452698 | | | | |
| TGF+SB42 | 1.207997 | 1.511181 | 0.223844 | 0.00173 | ** |
| | 1.778641 | | | | |
| | 1.394262 | | | | |
| | 1.663823 | | | | |
| TGF+5ITu 0.1µM | 1.948857 | 1.99059 | 0.188521 | 0.036942 | * |
| | 2.080224 | | | | |
| | 1.800191 | | | | |
| | 1.815342 | | | | |
| | 2.308334 | | | | |
| TGF+5ITu 1µM | 2.047267 | 1.843329 | 0.275074 | 0.012567 | * |
| | 1.800144 | | | | |
| | 2.262183 | | | | |

| | | | | | |
|---|---|---|---|---|---|
| | 1.374609 | | | | |
| | 1.719864 | | | | |
| | 1.855908 | | | | |
| TGF+5ITu 5µM | 1.150397 | 1.088089 | 0.161366 | 1.61E-05 | *** |
| | 0.838208 | | | | |
| | 1.039817 | | | | |
| | 1.335877 | | | | |
| | 1.076146 | | | | |
| TGF+5ITu 10µM | 0.430067 | 0.712802 | 0.150702 | 2.56E-06 | *** |
| | 0.808838 | | | | |
| | 0.915406 | | | | |
| | 0.715869 | | | | |
| | 0.649154 | | | | |
| | 0.75748 | | | | |

### Figure 2B

| ***SNAIL*** | **REP 1** | **REP 2** | **REP 3** | **AVERAGE** | **STDEV** | **P VALUE** | |
|---|---|---|---|---|---|---|---|
| CONTROL | 1 | 1 | 1 | 1 | 0 | 0.001292 | ** |
| TGF | 8.239786 | 8.327412 | 7.528274 | 8.031824 | 0.438283 | | |
| 5-ITu | 0.829903 | 0.720656 | 0.731161 | 0.760573 | 0.06027 | 0.001005 | ** |
| 5-ITu+TGF | 1.226427 | 0.879049 | 0.877526 | 0.994334 | 0.200999 | 0.000214 | *** |
| ABT | 1.066732 | 0.435222 | 1.005481 | 0.835812 | 0.34827 | 3.58E-05 | *** |
| ABT+TGF | 5.202591 | 5.887808 | 4.94672 | 5.345706 | 0.486594 | 0.00216 | ** |

| ***ZEB1*** | **REP 1** | **REP 2** | **REP 3** | **AVERAGE** | **STDEV** | **P VALUE** | |
|---|---|---|---|---|---|---|---|
| CONTROL | 1 | 1 | 1 | 1 | 0 | 0.022142 | * |
| TGF | 1.912496 | 2.570929 | 2.337449 | 2.273625 | 0.333824 | | |
| 5-ITu | 0.034088 | 0.051596 | 0.064718 | 0.050134 | 0.015367 | 0.007333 | ** |
| 5-ITu+TGF | 0.046352 | 0.060338 | 0.067559 | 0.058083 | 0.010782 | 0.007434 | ** |
| ABT | 0.630041 | 0.468887 | 0.980453 | 0.693127 | 0.261553 | 0.003585 | ** |
| ABT+TGF | 1.795072 | 2.161939 | 2.394972 | 2.117327 | 0.302428 | 0.580537 | ns |

### Figure 2C

| *SNAIL* | REP 1 | REP 2 | REP 3 | AVERAGE | STDEV | P VALUE | |
|---|---|---|---|---|---|---|---|
| CONTROL | 1 | 1 | 1 | 1 | 0 | 0.006832 | ** |
| TGF | 6.38933 | 6.501188 | 7.921935 | 6.937484 | 0.854392 | | |
| 5-ITu+TGF | 1.015345 | 0.081152 | 0.080581 | 0.392359 | 0.539521 | 0.000866 | *** |
| ABT+TGF | 6.716543 | 7.464375 | 9.070535 | 7.750484 | 1.202794 | 0.399319 | ns |

| *ZEB1* | REP 1 | REP 2 | REP 3 | AVERAGE | STDEV | P VALUE | |
|---|---|---|---|---|---|---|---|
| CTRL | 1 | 1 | 1 | 1 | 0 | 0.010759 | * |
| TGF | 1.472461 | 1.675161 | 1.548468 | 1.565363 | 0.102401 | | |
| 5-ITu+TGF | 0.282771 | 0.05459 | 0.052636 | 0.129999 | 0.132308 | 0.000176 | *** |
| ABT+TGF | 1.752743 | 2.403835 | 1.831856 | 1.996145 | 0.355279 | 0.162713 | ns |

### Figure 2F

**A549**

| | Migration index 18h | Average | Stdev | pvalue | |
|---|---|---|---|---|---|
| Control | 1.037055 | 1.225726 | 0.091027 | 5.57E-05 | *** |
| | 1.200113 | | | | |
| | 1.285462 | | | | |
| | 1.251892 | | | | |
| | 1.267909 | | | | |
| | 1.311922 | | | | |
| TGF | 1.617775 | 1.59137 | 0.080535 | | |
| | 1.501649 | | | | |
| | 1.661231 | | | | |
| | 1.618463 | | | | |
| | 1.684069 | | | | |
| | 1.465034 | | | | |
| TGF+5ITu 5µM | 0.911044 | 0.613682 | 0.292149 | 0.000811 | *** |
| | 0.399795 | | | | |
| | 0.567219 | | | | |
| | 0.641952 | | | | |

| | | | | | |
|---|---|---|---|---|---|
| | 0.548402 | | | | |
| TGF+ABT702 10 µM | 1.280658 | 1.261117 | 0.126958 | 0.00333 | ** |
| | 1.395525 | | | | |
| | 1.043015 | | | | |
| | 1.372307 | | | | |
| | 1.214078 | | | | |

### Figure 2G

**PANC1**

| Samples | Migration index 24h | Average | Stdev | pvalue | |
|---|---|---|---|---|---|
| Control | 1.340715 | 1.271352 | 0.041847 | 1.83E-09 | *** |
| | 1.275113 | | | | |
| | 1.274675 | | | | |
| | 1.255461 | | | | |
| | 1.210796 | | | | |
| TGF | 2.29046 | 2.249128 | 0.047993 | | |
| | 2.201489 | | | | |
| | 2.321921 | | | | |
| | 2.212033 | | | | |
| | 2.219735 | | | | |
| | 2.257148 | | | | |
| TGF+5ITu 5µM | 0.711044 | 0.713682 | 0.071629 | 6.86E-09 | *** |
| | 0.699795 | | | | |
| | 0.667219 | | | | |
| | 0.641952 | | | | |
| | 0.848402 | | | | |
| TGF+ABT702 10 µM | 2.180658 | 1.921117 | 0.285487 | 0.082532 | ns |
| | 1.595525 | | | | |
| | 1.643015 | | | | |
| | 1.872307 | | | | |
| | 2.314078 | | | | |

### Figure 3D

| Treatments | G1 (%) | | | S (%) | | | G2/M (%) | | | p-values (vs control) G2/M | S | G |
|---|---|---|---|---|---|---|---|---|---|---|---|---|
| control | 52.46 | 48.43 | 50.88 | 40.3 | 48.74 | 40.54 | 7.24 | 2.82 | 8.58 | | | |
| 5-ITu | 43.81 | 38.18 | 42.57 | 38.23 | 46.55 | 35.38 | 17.96 | 15.27 | 22.05 | 0.010057 | 0.511663 | 0.015519 |
| TGF | 97.6 | 74.53 | 64.32 | 2.08 | 25.4 | 35.54 | 0.33 | 0.07 | 0.13 | 0.073702 | 0.146338 | 0.101121 |
| TGF+5-ITu | 44.5 | 43.9 | 39.24 | 35.88 | 41.78 | 35.77 | 19.62 | 14.32 | 24.99 | 0.029314 | 0.196984 | 0.020579 |

### Figure 3E

| **A549** | **Rep 1** | **Rep 2** | **Rep 3** | **Avge** | **SD** | *p-value* | |
|---|---|---|---|---|---|---|---|
| Control | 4735 | 4767 | 5051 | 4751 | 16 | 0.001381 | ** |
| TNF | 4938 | 5190 | 5321 | 5064 | 126 | 0.001325 | ** |
| TNF+5-ITu | 9528 | 10246 | 10527 | 9887 | 359 | | |
| TNF+5-ITu+Nec-1 | 6485 | 7773 | 7620 | 7129 | 644 | 0.006538 | ** |

| **PANC1** | Rep 1 | Rep 2 | Rep 3 | **Avge** | **SD** | *p-value* | |
|---|---|---|---|---|---|---|---|
| Control | 7744 | 6254 | 7091 | 6999 | 745 | 0.000146 | *** |
| TNF | 7717 | 8768 | 8163 | 8242.5 | 525.5 | 0.000476 | *** |
| TNF+5-ITu | 25046 | 26766 | 27659 | 25906 | 860 | | |
| TNF+5-ITu+Nec-1 | 11132 | 10617 | 11102 | 10874.5 | 257.5 | 0.001642 | ** |

### Figure 3F

| ***CDH2*** | **REP1** | **REP2** | **REP3** | **AVERAGE** | **STDEV** | **p value** | |
|---|---|---|---|---|---|---|---|
| Control | 1 | 1 | 1 | 1 | 0 | 0.002452 | ** |
| TGFβ | 6.152517 | 5.466996 | 6.247317 | 5.95561 | 0.347663 | | |
| TGFβ+ITU | 1.309668 | 1.344606 | 1.343706 | 1.33266 | 0.016262 | 0.002766 | ** |
| TGFβ+ITU+Nec1 | 0.949957 | 0.84868 | 0.925036 | 0.907891 | 0.043087 | 0.002076 | ** |
| TGFβ+ITU+GSK | 1.498935 | 1.579312 | 1.662076 | 1.580108 | 0.066604 | 0.002362 | ** |

| *SNAI1* | REP1 | REP2 | REP3 | AVERAGE | STDEV | p value | |
|---|---|---|---|---|---|---|---|
| Control | 1 | 1 | 1 | 1 | 0 | 0.000362 | *** |
| TGFβ | 9.576292 | 9.029273 | 9.287278 | 9.297614 | 0.223439 | | |
| TGFβ+ITU | 0.303595 | 0.349897 | 0.305053 | 0.319515 | 0.021492 | 0.000276 | *** |
| TGFβ+ITU+Nec1 | 0.262099 | 0.285766 | 0.2903 | 0.279388 | 0.012365 | 0.000295 | *** |
| TGFβ+ITU+GSK (5μM) | 0.341367 | 0.407449 | 0.36733 | 0.372049 | 0.027183 | 0.00026 | *** |

**Figure 4C**

| *ZEB1* | Rep 1 | Rep 2 | Rep 3 | AVG | STDEV | p value | |
|---|---|---|---|---|---|---|---|
| Control | 1 | 1 | 1 | 1 | 0 | 0.010263237 | * |
| TGF | 6.76347 | 8.50318 | 6.45681 | 7.24116 | 1.10365 | | |
| BMS+TGF | 0.18046 | 0.39708 | 0.16669 | 0.24808 | 0.12923 | 0.00755502 | ** |
| TGF+5-ITu | 0.05534 | 0.1123 | 0.0403 | 0.06931 | 0.03798 | 0.007745004 | ** |

| *SNAI1* | Rep 1 | Rep 2 | Rep 3 | AVG | STDEV | p value | |
|---|---|---|---|---|---|---|---|
| Control | 1 | 1 | 1 | 1 | 0 | 0.008412743 | ** |
| TGF | 6.74204 | 8.55475 | 8.80074 | 8.03251 | 1.12433 | | |
| BMS+TGF | 0.90341 | 0.71148 | 0.41208 | 0.67565 | 0.24762 | 0.005764756 | ** |
| TGF+5-ITu | 0.73362 | 0.55528 | 0.38116 | 0.55669 | 0.17624 | 0.006414436 | ** |

| *CDH2* | Rep 1 | Rep 2 | Rep 3 | AVG | STDEV | *p value* | |
|---|---|---|---|---|---|---|---|
| Control | 1 | 1 | 1 | 1 | 0 | 0.015758066 | * |
| TGF | 5.21728 | 7.51198 | 6.07695 | 6.26874 | 1.15931 | | |
| BMS+TGF | 0.50112 | 0.63656 | 0.4392 | 0.52563 | 0.10093 | 0.012810955 | * |
| TGF+5-ITu | 0.44089 | 0.5447 | 0.42597 | 0.47052 | 0.06468 | 0.012858727 | * |

| *VIM* | Rep 1 | Rep 2 | Rep 3 | AVG | STDEV | *p value* | |
|---|---|---|---|---|---|---|---|
| Control | 1 | 1 | 1 | 1 | 0 | 0.043557922 | * |
| TGF | 10.67477 | 6.622598 | 13.51972 | 10.27236 | 3.466127 | | |
| BMS+TGF | 1.706958 | 1.313741 | 1.592654 | 1.537784 | 0.202269 | 0.048250214 | * |
| ITU+TGF | 0.86737 | 0.937917 | 0.813634 | 0.872974 | 0.062331 | 0.042421994 | * |

### Figure 4E

| | Rep 1 | Rep 2 | Rep 3 | Average | stdev | *p value (TNF)* | *p value (TGF)* |
|---|---|---|---|---|---|---|---|
| **Control** | 1.10687 | 1.384793 | 1.382151 | 1.291271 | 0.130396 | 0.006709658 | 0.000828 |
| **TNF** | 59.72622 | 46.8341 | 47.85412 | 51.47148 | 5.85182 | | |
| **TNF+BMS345** | 3.619632 | 4.502959 | 4.032836 | 4.051809 | 0.360866 | 0.007353447 | |
| **TNF+5-ITu** | 2.723837 | 3.044944 | 2.39823 | 2.722337 | 0.264022 | 0.007036951 | |
| **TGF** | 6.529268 | 7.403061 | 6.735426 | 6.889252 | 0.372939 | 0.000382231 | |
| **TGF+BMS345** | 4.304183 | 3.398964 | 3.73494 | 3.812695 | 0.373622 | | 0.001182 |
| **TGF+5-ITu** | 2.537433 | 2.591647 | 2.755611 | 2.62823 | 0.092751 | | 0.002428 |

### Figure 5C

| ***SMAD3*** | **Rep1** | **Rep2** | **Rep3** | **Average** | **St dev** | ***p value*** | |
|---|---|---|---|---|---|---|---|
| Control | 1 | 1 | 1 | 1 | 0 | 0.01903 | * |
| TGF | 2.214958 | 2.199633 | 2.796508 | 2.4037 | 0.277828 | | |
| TGF+5ITu | 0.08874 | 0.131529 | 0.069453 | 0.096574 | 0.025941 | 0.006792 | ** |
| ***SERPINE1*** | **Rep1** | **Rep2** | **Rep3** | **Average** | **St dev** | ***p value*** | |
| Control | 1 | 1 | 1 | 1 | 0 | 0.019387 | * |
| TGF | 21.10236 | 33.49367 | 25.4584 | 26.68481 | 5.132526 | | |
| TGF+5ITu | 0.128643 | 0.650691 | 0.133726 | 0.304353 | 0.244907 | 0.01822 | * |
| ***TGIF*** | **Rep1** | **Rep2** | **Rep3** | **Average** | **St dev** | ***p value*** | |
| Control | 1 | 1 | 1 | 1 | 0 | 0.024094 | * |
| TGF | 3.478652 | 2.476313 | 2.739064 | 2.898009 | 0.424357 | | |
| TGF+5ITu | 0.300676 | 0.194507 | 0.203349 | 0.232844 | 0.0481 | 0.01164 | * |
| ***PMEPA1*** | **Rep1** | **Rep2** | **Rep3** | **Average** | **St dev** | ***p value*** | |
| Control | 1 | 1 | 1 | 1 | 0 | 0.005879 | ** |
| TGF | 9.282689 | 11.26304 | 9.140487 | 9.895405 | 0.968804 | | |
| TGF+5-ITu | 0.225265 | 0.719744 | 0.243313 | 0.396108 | 0.228964 | 0.003548 | ** |

### Figure 5D

| *JUNB* | Rep1 | Rep2 | Rep3 | Average | St dev | *p value* | |
|---|---|---|---|---|---|---|---|
| Control | 1 | 1 | 1 | 1 | 0 | 0.137284 | ns |
| TGF | 73.39635 | 13.0172 | 42.7497 | 43.05442 | 24.65063 | | |
| TGF+5ITu | 785.0371 | 62.31036 | 449.1019 | 432.1498 | 295.2954 | 0.202695 | ns |
| *ID2* | Rep1 | Rep2 | Rep3 | Average | St dev | *p value* | |
| Control | 1 | 1 | 1 | 1 | 0 | 0.624696 | ns |
| TGF | 1.284334 | 0.412949 | 0.870416 | 0.8559 | 0.35589 | | |
| TGF+5-ITu | 1.129373 | 0.902196 | 0.863526 | 0.965031 | 0.117275 | 0.713998 | ns |
| *PPP1R15A* | Rep1 | Rep2 | Rep3 | Average | St dev | *p value* | |
| Control | 1 | 1 | 1 | 1 | 0 | 0.010496 | * |
| TGF | 0.683183 | 0.689628 | 0.774303 | 0.715705 | 0.041519 | | |
| TGF+5-ITu | 0.857109 | 1.362205 | 0.966175 | 1.061829 | 0.217014 | 0.148396 | ns |

### Figure 5E

| *XIAP* | Rep1 | Rep2 | Rep3 | Average | St dev | *p value* | |
|---|---|---|---|---|---|---|---|
| Control | 1 | 1 | 1 | 1 | 0 | 0.040649 | * |
| TGF | 1.454363 | 1.981378 | 1.776336 | 1.737359 | 0.216911 | | |
| TGF+5ITu | 0.328707 | 0.122133 | 0.353164 | 0.268001 | 0.103627 | 0.003865 | ** |

### Figure S1B

| *SNAIL* | Rep 1 | Rep 2 | Rep 3 | AVG | STDEV | *p-value* | |
|---|---|---|---|---|---|---|---|
| Control | 0.115771 | 0.089817 | 0.096511 | 0.100699 | 0.013475 | 7.48E-05 | *** |
| TGF | 1 | 1 | 1 | 1 | 0 | | |
| SB42 | 0.107551 | 0.080673 | 0.098809 | 0.095678 | 0.01371 | 7.66E-05 | *** |
| SB42+TGF | 0.115577 | 0.094407 | 0.119121 | 0.109702 | 0.013364 | 7.51E-05 | *** |

| | | | | | | | |
|---|---|---|---|---|---|---|---|
| ITU 5 um | 0.186251 | 0.22699 | 0.189125 | 0.200789 | 0.022737 | 0.00027 | *** |
| ITU+TGF 5 um | 0.199031 | 0.180508 | 0.211403 | 0.196981 | 0.015549 | 0.000125 | *** |

| ***ZEB1*** | **Rep 1** | **Rep 2** | **Rep 3** | **AVG** | **STDEV** | ***p-value*** | |
|---|---|---|---|---|---|---|---|
| Control | 0.693842 | 0.61122 | 0.514594 | 0.606552 | 0.089715 | 0.016894 | * |
| TGF | 1 | 1 | 1 | 1 | 0 | | |
| SB42 | 0.639608 | 0.643736 | 0.657101 | 0.646815 | 0.009144 | 0.000223 | *** |
| SB42+TGF | 0.701953 | 0.730711 | 0.710152 | 0.714272 | 0.014815 | 0.000895 | *** |
| ITU 5 um | 0.300449 | 0.301669 | 0.252903 | 0.285007 | 0.02781 | 0.000504 | *** |
| ITU+TGF 5 um | 0.342571 | 0.390814 | 0.119283 | 0.284223 | 0.144864 | 0.01338 | * |

### Figure S3A

| ***SNAI1*** | **Rep 1** | **Rep 2** | **Rep 3** | **AVG** | **STDEV** | ***p-value*** | |
|---|---|---|---|---|---|---|---|
| CTRL | 1 | 1 | 1 | 1 | 0 | 0.00683 | ** |
| TGF | 6.38933 | 6.50119 | 7.92193 | 6.93748 | 0.85439 | | |
| BMS+TGF | 1.72789 | 1.63154 | 0.72428 | 1.36124 | 0.55372 | 0.00141 | ** |
| ITU+TGF | 1.01534 | 0.08115 | 0.08058 | 0.39236 | 0.53952 | 0.00087 | *** |
| ABT+TGF | 6.71654 | 7.46438 | 9.07053 | 7.75048 | 1.20279 | 0.39932 | ns |

| ***ZEB1*** | **Rep 1** | **Rep 2** | **Rep 3** | **AVG** | **STDEV** | ***p-value*** | |
|---|---|---|---|---|---|---|---|
| CTRL | 1 | 1 | 1 | 1 | 0 | 0.01076 | * |
| TGF | 1.47246 | 1.67516 | 1.54847 | 1.56536 | 0.1024 | | |
| BMS+TGF | 0.85201 | 1.10815 | 0.83171 | 0.93063 | 0.15407 | 0.51709 | ns |
| ITU+TGF | 0.28277 | 0.05459 | 0.05264 | 0.13 | 0.13231 | 0.00762 | ** |
| ABT+TGF | 1.75274 | 2.40384 | 1.83186 | 1.99614 | 0.35528 | 0.16271 | ns |

### Figure S3C

| Reaction | Rep 1 (%) | Rep 2 (%) | Rep 3 (%) | AVERAGE | STDEV | *p-value* | |
|---|---|---|---|---|---|---|---|
| IKBA | 0 | 0 | 0 | 0 | | NA | |
| IKK+IKBA | 100 | 100 | 100 | 100 | 0 | | |
| IKK+IKBA+5-Itu 0.1μM | 41.8070 | 45.2898 | 22.7064 | 36.6011 | 9.9274 | 0.0120 | * |
| IKK+IKBA+5-Itu 1μM | 28.7715 | 33.9868 | 17.4468 | 26.7350 | 6.9043 | 0.0044 | ** |
| IKK+IKBA+5-Itu 5μM | 15.0378 | 22.0303 | 12.0828 | 16.3836 | 4.1711 | 0.0012 | ** |
| IKK+IKBA+BMS 0.1μM | 24.8585 | 39.9041 | 35.8701 | 33.5443 | 6.3587 | 0.0045 | ** |
| IKK+IKBA+BMS 1μM | 3.3974 | 38.2999 | 19.9386 | 20.5453 | 14.2553 | 0.0157 | * |
| IKK+IKBA+BMS 5μM | 4.5200 | 28.1856 | 13.1476 | 15.2844 | 9.7789 | 0.0066 | ** |
