## Supplementary Table S1 for "5-Iodotubercidin inhibits Epithelial to Mesenchymal Transition by inhibiting IKK/NFκB-dependent gene expression"

| Supplementary Table S1: Oligonucleotide sequences used for real time qPCR | | | |
|---|---|---|---|
| **Sl.No** | **Names** | **species** | **Primer Sequences** |
| 1 | hSMAD3-Fwd | Human | TCGAGCCCCAGAGCAATATT |
| 2 | hSMAD3-Rev | Human | CGTCCATGCTGTGGTTCATC |
| 3 | SERPINE1-Fwd | Human | TCGAGGTGAACGAGAGTGGCA |
| 4 | SERPINE1-Rev | Human | AAGGACTGTTCCTGTGGGGTTGT |
| 5 | hTGIF1-Fwd | Human | GACATTCCCTTGGACCTT TCT |
| 6 | Htgif1-Rev | Human | TACAGCCAATCCCGAAGAATC |
| 7 | Hpmepa1-Fwd | Human | TGTCAGGCAACGGAATCCC |
| 8 | hPMEPA1-Rev | Human | CAGGTACGGATAGGTGGGC |
| 9 | hID2-Fwd | Human | ATGAAAGCCTTCAGTCCGGTG |
| 10 | hID2-Rev | Human | AGCAGACTCATCGGGTCGT |
| 11 | hPPP1R15A-Fwd | Human | GAATCAAGCCACGGAGGATA |
| 12 | hPPP1R15A-Rev | Human | CAGGGAGGACACTCAGCTTC |
| 13 | hJUNB-Fwd | Human | CGACCACCATCAGCTACCTC |
| 14 | Hjunb-Rev | Human | GTCTGCGGTTCCTCCTTGAA |
| 15 | hXIAP-Fwd | Human | TGGTATCCAGGGTGCAAATATCT |
| 16 | hXIAP-Rev | Human | GTAGTTCTTACCAGACACTCCTCA |
| 17 | hKLF10_fwd | Human | GCCAACCATGCTCAACTTCG |
| 18 | hKLF10_rev | Human | TGCAGTTTTGTTCCAGGAATACAT |
| 19 | hBMP2_fwd | Human | ACCCGCTGTCTTCTAGTGTTG |
| 20 | hBMP2_rev | Human | TTCTTCGTGATGGAAGCTGAG |
| 21 | hIFNGR2 Fwd | Human | GCAAGATTCGCCTGTACAACG |
| 22 | hIFNGR2 Rev | Human | GTCACCTCAATCTTTTCTGGAG |
| 23 | huCDH2_qPCR_F | Human | TGAGGAGTCAGTGAAGGAGTCA |
| 24 | huCDH2_qPCR_R | Human | CCAGTCTCTCTTCTGCCTTTGT |
| 25 | huCDH1_qPCR_F | Human | CCCGGGACAACGTTTATTAC |
| 26 | huCDH1_qPCR_R | Human | GCTGGCTCAAGTCAAAGTCC |
| 27 | sy-huSNAI1/122s | Human | TCGGAAGCCTAACTACAGCG |
| 28 | sy-huSNAI1/277as | Human | GCCAGGACAGAGTCCCAGA |
| 29 | huCyclophilin_qPCR_F | Human | GTCAACCCCACCGTGTTCTT |
| 30 | huCyclophilin_qPCR_R | Human | CTGCTGTCTTTGGGACCTTGT |
| 31 | huVIM_qPCR_F | Human | TGAGTACCGGAGACAGGTGCAG |
| 32 | huVIM_qPCR_R | Human | TAGCAGCTTCAACGGCAAAGTTC |
| 33 | sy-huZEB-1/s | Human | TTTCCCATTCTGGCTCCTAT |
| 34 | sy-huZEB-1/as | Human | GATGCTGAAAGAGACGGTGA |
| 35 | hu GAPDH_Fwd | Human | TGCACCACCAACTGCTTAGC |
| 36 | hu GAPDH_Rev | Human | GGCATGGACTGTGGTCATGAG |
